# Probiotic bacteria and bile acid profile are modulated by prebiotic diet and associate with facilitated diurnal clock/sleep realignment after chronic disruption of rhythms

**DOI:** 10.1101/2021.03.03.433775

**Authors:** Robert S. Thompson, Michelle Gaffney, Shelby Hopkins, Tel Kelley, Antonio Gonzalez, Samuel J. Bowers, Martha Hotz Vitaterna, Fred W. Turek, Christine L. Foxx, Christopher A. Lowry, Fernando Vargas, Pieter C. Dorrestein, Kenneth P. Wright, Rob Knight, Monika Fleshner

## Abstract

Chronic disruption of rhythms (CDR) impacts sleep and can result in circadian misalignment of physiological systems, which in turn is associated with increased disease risk. Exposure to repeated or severe stressors also disturbs sleep and diurnal rhythms. Prebiotic nutrients produce favorable changes in gut microbial ecology, the gut metabolome, and reduce several negative impacts of acute severe stressor exposure, including disturbed sleep, core body temperature rhythmicity, and gut microbial dysbiosis. This study tested the hypothesis whether prebiotics can also reduce the negative impacts of CDR by facilitating light/dark realignment of sleep/wake, core body temperature, and locomotor activity; and whether prebiotic-induced changes in bacteria and bile acid profiles are associated with these effects. Male, Sprague Dawley rats were fed diets enriched in prebiotic substrates or calorically matched control chow. After 5 weeks on diet, rats were exposed to CDR (12h light/dark reversal, weekly for 8 weeks) or remained on undisturbed normal light/dark cycles (NLD). Sleep EEG, core body temperature, and locomotor activity were recorded via biotelemetry in freely moving rats. Fecal samples were collected on experimental days -33, 0 (day of onset of CDR), and 42. Taxonomic identification and relative abundances of gut microbes were measured in fecal samples using 16S rRNA gene sequencing and shotgun metagenomics. Fecal primary, bacterially-modified secondary, and conjugated bile acids were measured using liquid chromatography with tandem mass spectrometry (LC-MS/MS). Prebiotic diet produced rapid and stable increases in the relative abundances of *Parabacteroides distasonis* and *Ruminiclostridium 5*. Shotgun metagenomics analyses confirmed reliable increases in relative abundances of *Parabacteroides distasonis* and *Clostridium leptum*, a member of the *Ruminiclostridium* genus. Prebiotic diet also modified fecal bile acid profiles; and based on correlational and step-wise regression analyses, *Parabacteroides distasonis* and *Ruminiclostridium 5* were positively associated with each other and negatively associated with secondary and conjugated bile acids. Prebiotic diet, but not CDR, impacted beta diversity. Measures of alpha diversity evenness were decreased by CDR and prebiotic diet prevented that effect. Rats exposed to CDR while eating prebiotic, compared to control diet, more quickly realigned NREM sleep and core body temperature (ClockLab) diurnal rhythms to the altered light/dark cycle. Finally, both cholic acid and *Ruminiclostridium 5* prior to CDR were associated with time to realign CBT rhythms to the new light/dark cycle after CDR; whereas both *Ruminiclostridium 5* and taurocholic acid prior to CDR were associated with NREM sleep recovery after CDR. These results suggest that ingestion of prebiotic substrates is an effective strategy to increase the relative abundance of health promoting microbes, alter the fecal bile acid profile, and facilitate the recovery and realignment of sleep and diurnal rhythms after circadian disruption.

## 1. Introduction

Most biological organisms oscillate in circadian cycles of approximately 24-hour periods in coordination with the light/dark cycle. The coordination and regulation of circadian rhythms requires complex integration of brain-controlled signals, molecular gene clocks, as well as environmental cues (Vitaterna, Shimomura, and Jiang 2019). The light/dark cycle and diet (Voigt et al. 2016) are potent environmental cues that influence circadian rhythms. External light/dark cues help entrain the master pacemaker in the brain, the suprachiasmatic nucleus, while a consistent schedule of food ingestion (Thaiss et al. 2014) and nutrient composition (Kohsaka et al. 2007; Tahara et al. 2018) function as entrainment cues to peripheral clocks in the liver and intestine (Oosterman et al. 2015). Ideally, these entrainment cues (central and peripheral) and circadian clocks throughout the body are synchronized to support optimal function.

Rapidly changing entrainment cues that are misaligned or in opposition of previously aligned biological rhythms produces physiological strain on the organism resulting in negative health consequences (Depner, Stothard, and Wright 2014; Marcheva et al. 2010; Turek et al. 2005; Morris et al. 2016). Members of the military, first responders, pilots, astronauts, and health care professionals frequently experience chronic disruption of rhythms (CDR) due to altered light exposure, repeated jet lag, disturbed/restricted sleep, and eating unhealthy food at the wrong biological times of day. Unfortunately, CDR and circadian misalignment of physiological rhythms are common for many members of society. There is, therefore, a need for effective, non-invasive, readily translatable ways of facilitating realignment and reducing the probability of physiological rhythm misalignment. We propose that consumption of select dietary substrates could be a novel intervention for reducing the physiological strain of CDR.

Prebiotics are dietary substrates that are selectively utilized by host microorganisms conferring health benefits (Gibson et al. 2017; Marco et al. 2021). Prebiotic substrates: 1) resist gastric acidity, hydrolysis by mammalian enzymes, and absorption in the upper gastrointestinal tract; 2) are fermented by intestinal commensal bacteria; and 3) selectively stimulate the growth and/or activity of intestinal bacteria associated with health and well-being (Klosterbuer, Roughead, and Slavin 2011; Koecher, Thomas, and Slavin 2015; Brownawell et al. 2012; Gu et al. 2019). Health benefits produced by prebiotics include reductions in inappropriate inflammation (Chang et al. 2018), intestinal dysbiosis (Cai et al. 2018), sleep/wake disturbances (Thompson et al. 2017), and anxiety-like behavior (Mika et al. 2018; Tarr et al. 2015). There is mounting evidence that dietary prebiotics impact the diversity and composition of intestinal bacterial communities, however many studies are limited to phyla level changes measured at a single time point (Cai et al. 2018; Xie et al. 2018; Ferrario et al. 2017). There is a need, therefore, for additional prebiotic studies that assess changes in the diversity and composition of intestinal bacterial communities with greater taxonomic and chronological resolution.

Galactooligosaccharide (GOS) and polydextrose (PDX) are examples of dietary prebiotics that have been reported to produce favorable changes in gut microbial ecology, to alter the relative abundance of bacterially derived metabolites and to promote a stress robust phenotype (Mika et al. 2018; Thompson et al. 2020; Mika et al. 2016). We and others have evidence from preclinical acute stress studies that GOS/PDX reduces anxiety-like behavior (Mika et al. 2016) and alleviates stress-induced disruption of diurnal core body temperature and sleep (Thompson et al. 2017). GOS/PDX produces phyla-level changes in gut microbes that are associated with increased non-rapid eye movement (NREM) sleep and attenuates the negative impact of stress on fecal microbial alpha diversity (Thompson et al. 2017). Finally, GOS/PDX can alter neurobiologically active microbially derived gut metabolites in response to stressor exposure (Thompson et al. 2020). Taken together these studies suggest that GOS/PDX can alter the gut microbial ecology and significantly reduce the impacts of acute stressor exposure on host physiological variables like core body temperature and sleep. It remains unknown, however, if GOS/PDX can reduce the physiological strain of CDR and/or impact the rate of circadian realignment of host physiological rhythms like core body temperature, locomotor activity, or sleep.

The mechanisms for how prebiotic-induced changes in gut microbial ecology impact complex brain functions and modulate physiology are largely unknown. There is evidence that changes in microbially derived secondary bile acid profiles, for example, could play a role (McMillin and DeMorrow 2016; An, Zhao, and Liu 2019; Perino et al. 2020). Bile acid enterohepatic cycling is a complex and dynamic process (Mertens et al. 2017). In general, primary bile acids are synthesized in the liver from cholesterol and released into the intestine after food intake. The dominant primary bile acids are cholic acid (CA) and chenodeoxycholic acid (CDCA) in humans; and CA and beta-muricholic acid (β−MCA) in rodents. To facilitate bile acid transport from the liver and absorption in intestine, primary bile acids are glycine or taurine conjugated to produce conjugated bile acids such as taurocholic acid (TCA), tauroursodeoxycholic acid (TUDCA), or glycourosodeoxycholic (GUDCA). Once in the colon, conjugated bile acids undergo microbially mediated transformations to produce secondary and secondary conjugated bile acids such as lithocholic acid (LCA), deoxycholic acid (DCA), hyodeoxycholic acid (HDCA), and GUDCA. Approximately 95% of the bile acids are reabsorbed in the intestine and/or released into the portal vein and ∼5% is excreted in feces. Changes in relative levels or concentration of fecal bile acids could reflect differences in microbial biotransformation, re-absorption, and/or primary bile acid synthesis. Enterohepatic cycling and bile acid synthesis are under circadian control (Ho 1976; Yu et al. 2020; Ma et al. 2009) and are regulated by bile acid sensors, farnesoid X receptor (FXR) and Takeda G protein-coupled receptor 5 (TGR5). These receptors are found throughout the body and brain, with dense expression in the intestine (Al-Aqil et al. 2018; Chen, Yang, et al. 2017; Chiang et al. 2017). Primary and secondary bile acids can bind both FXR and TGR5 with varying affinities, and act as receptor agonists or antagonists (Sepe et al. 2016). It is feasible; therefore, that dietary prebiotics impact the brain and physiology by changing microbially mediated bile acid composition and bile acid signaling.

The current study tested the hypothesis whether prebiotics can reduce the negative impacts of CDR by facilitating light/dark realignment of sleep/wake, core body temperature, and locomotor activity rhythms; and potential health promoting gut microbial changes and bile acid profiles are associated with these effects.

## 2. Methods

### 2.1 Animals

Adult male Sprague Dawley rats (n = 84, Harlan Laboratories, Indianapolis, IN) were housed with controlled temperature and humidity. All procedures were approved by the University of Colorado Institutional Animal Care and Use Committee as previously described (Thompson et al. 2017; Thompson et al. 2020). Briefly, animals weighed 40-50g upon arrival at post-natal day (PND) 23 and were maintained on a 12:12 h light/dark cycle. All rats were housed in Nalgene Plexiglas cages and were placed on control or GOS/PDX diet *(ad libitum)* on arrival. This study is a part of a multi-site, multi-species, Department of Defense and Office of Naval Research-funded project designed to understand the impacts of chronic disruption of rhythms common to many in the military. Given this population of the Navy is ∼80-90% male and to not exceed the award budget, only male rats were tested in this study.

### 2.2 Experimental Design

The detailed experimental design is depicted in Figure 1. Rats arrived on experimental day -35 and were placed on either control (n=42) or prebiotic diet (n=42, GOS/PDX) for the duration of the study. To optimize housing acclimatization, rats were double housed and remained in their home cages undisturbed until experimental day -7 when all animals were instrumented with F40-EET biotelemetry devices. After surgery, all rats were single housed for the remainder of the study as required for *in vivo* biotelemetry. Chronic disruption of rhythms (CDR, n = 42) was induced by a 12-hour reversal of the light/dark cycle each week for 8 weeks. Control animals’ light/dark cycles were undisturbed and consistent with the normal light/dark cycle (NLD, n = 42) of the vivarium. Continuous biotelemetry signals were collected from experimental day -1 until the end of the experiment on day 56; diurnal rhythm analyses of core body temperature and locomotor activity were used to document light/dark rhythm misalignment and realignment.

**FIG. 1.**
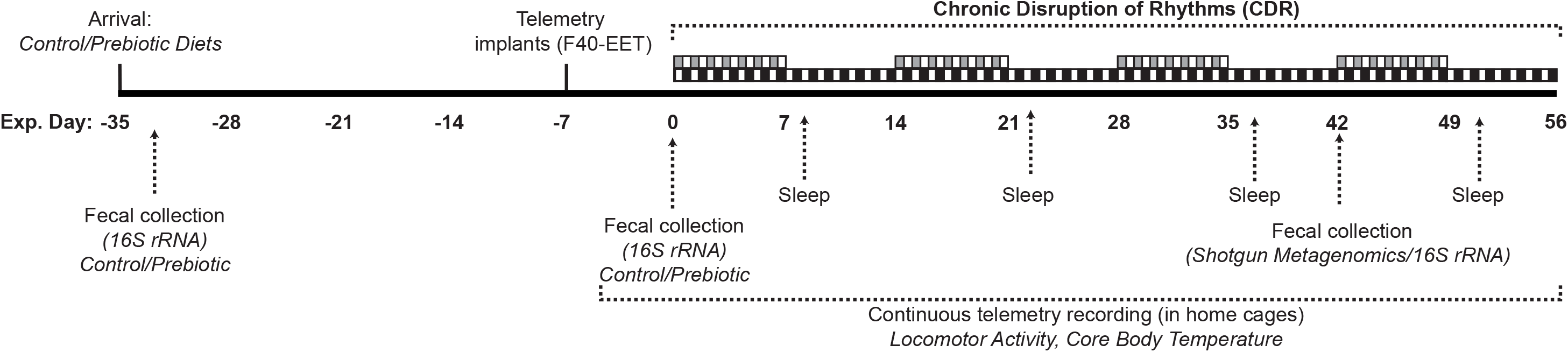
Experimental schematic detailing the experimental timeline. Rats (postnatal day 23) arrived on experimental day -35 and were placed on either control or prebiotic diet (GOS/PDX) for the duration of the study. Rats remained in their home cages undisturbed until experimental day -7 when all animals were instrumented with F40-EET biotelemetry devices. Chronic disruption of rhythms (CDR) was induced by reversing the light/dark cycle 12-hours each week for 8 weeks. Control animals’ light/dark cycles were undisturbed and consistent with the normal light/dark cycle (NLD) of the vivarium. Continuous biotelemetry signals were collected from experimental day -1 until the end of the experiment on day 56. Thirty-six-hour samples of electroencephalogram (EEG) data were collected starting on experimental days 8, 22, 36, and 50 were used to analyze changes in sleep architecture due to CDR and diet. To capture the time course of prebiotic-induced changes and the impact of a history of CDR on the microbiome and bile acids profiles, fecal samples were collected on experimental days -33, 0, and 42.

### 2.3 Diets

Diet contained the following prebiotic substrates, which were absent from control diet: galactooligosaccharides (GOS, 24.14g/Kg (7.0 g active); FrieslandCampina, Zwolle, The Netherlands), polydextrose (PDX, 7.69g/Kg (7.0 g active); Danisco, Terre Haute, IN, USA). GOS is a non-absorbable complex carbohydrate derived from the enzymatic breakdown of lactose and is enriched in whole grains, lentils, beans, and cabbage, while PDX is a processed polymer derived from glucose and classified as a soluble fiber. GOS/PDX are prebiotic nutrients digested by commensal gut bacteria and are thought to increase the abundance of probiotic bacterial species such as *Lactobacillus* spp. (Cardelle-Cobas et al. 2011; Herfel et al. 2011; Saulnier et al. 2013). The diets were originally formulated by Mead Johnson Nutrition (MJN, Evansville, IN, USA) based on AIN-93G specifications and were isocaloric with similar carbohydrate, protein, fat, vitamin, and mineral levels details of which have been previously published (Mika et al. 2016; Thompson et al. 2017; Thompson et al. 2020).

### 2.4 Fecal Sample Collection Procedures

Weekly fecal samples were collected in the light cycle shortly after cage changes (∼1100 hrs) in order to minimize disruption to the animals. Sterile forceps (100% ethanol) were used to collect fecal samples, placed into 1.5 mL sterile airtight screw caps tubes (USA Scientific, Ocala, FL, USA), and then submerged in liquid nitrogen for rapid freezing.

### 2.5 Biotelemetric Surgery

Biotelemetry surgeries were performed as previously described in detail (Thompson et al. 2013; Thompson et al. 2016; Thompson et al. 2017; Thompson et al. 2014; Belfry et al. 2012), with the exception of *a priori* elimination of antibiotic administration post-operation, in order to eliminate this as a potential confounding variable on the gut microbiome.

### 2.6 Biotelemetric Data Acquisition

Data acquisition/analysis was as described previously in detail (Belfry et al. 2012; Greenwood et al. 2014; Thompson et al. 2013; Thompson et al. 2016). F40-EET transmitter allows *in vivo* real-time measurement of locomotor activity (LA), core body temperature (CBT), and electroencephalogram (EEG) in freely behaving animals. Biotelemetry recordings were acquired using Dataquest ART 4.3 Gold Acquisition Software (Data Sciences International, St. Paul, MN).

### 2.7 Circadian Analysis of Host Physiology

Biotelemetry recordings of CBT and LA were analyzed using Dataquest ART 4.3 Gold Analysis Software (Data Sciences International, St. Paul, MN) and circadian cycles were scored using the ClockLab program version 6.1.02 (Actimetrics Software, Wilmette, IL, USA). Clocklab analysis was used to mark, examine, and measure the bathyphase (or minimum) of core body temperature across days. The bathyphase is used as a marker of circadian timing (Benloucif et al. 2008). The bathyphase of locomotor activity was also measured and quantified. The bathyphase is automatically calculated in the Clocklab analysis software and is based on sine wave approximation of 5-min interval data. Once all bathyphase data were marked, the number of hours to realign was counted by an observer blind to treatment of each animal. Each animal’s recording was used as its own control since each rodent is compared to its own intrinsic clock rhythm (i.e., bathyphase). The maximum number of hours was 168 each week (7 days x 24 hours). A failure to realign to the new original light/dark cycle occurred if after 168 hours the rat was not back in alignment with its own bathyphase.

### 2.8 Biotelemetric Sleep Analyses

Analysis of the sleep/wake cycles was performed using the automated Neuroscore 3.1.0 software (Data Sciences International). The trace EEG signal was subjected to fast Fourier Transformation (FFT), yielding spectra between 0.5 and 30 Hz in 0.5-Hz frequency bins. The delta frequency band was defined at 0.5-4.5 Hz and the theta frequency band was defined as 6.0-9.0 Hz as previously described (Olivadoti and Opp 2008; Thompson et al. 2016). Sleep recordings were autoscored and then corrected by an observer blind to the experimental treatment of each animal. Each recording was autoscored/corrected in 10-sec epochs and classified as NREM, REM, or WAKE on the basis of state-dependent changes in multiple parameters, including the EEG, locomotor activity, and CBT as previously described (Thompson et al. 2016; Thompson et al. 2017).

#### 2.8.1 Rationale for Sleep Snapshot Analyses

Electroencephalogram (EEG) sleep data were collected on experimental days 8, 22, 36, and 50, and were used to analyze changes in sleep architecture due to CDR and diet. Thirty-six hours of sleep snapshots across the duration of the study were targeted for the following reasons: 1) this snapshot captures the early process of realignment of sleep/wake states to the new light/dark cycle; 2) the sleep/wake states for both NLD and CDR groups are measured at matched clock times; and 3) analyses of sleep snapshots after each of the 4 cycles of repeated circadian disruption will reveal potential cumulative effects of chronic disruption of rhythms.

Time spent in NREM/REM/WAKE was calculated as a percentage (%) of time spent in a specific behavioral state per hour (%). Average bout duration per hour of NREM/REM/WAKE was also calculated. Bout duration is defined as the duration of time from state change onset to state change offset with a temporal resolution of 10 seconds. Total number of episodes per hour (#) of NREM/REM/WAKE was also calculated. An episode is an identified change in sleep/wake state with a temporal resolution of 10 seconds. All sleep/wake parameters were averaged into 12-hour bins for statistical analysis.

### 2.9 Fecal Microbiome

Fecal samples for 16S rRNA gene analysis were collected on experimental day -33, 0 and 42. Samples were collected and prepared as previously described (Mika and Fleshner 2016; Maslanik et al. 2013). Briefly, after purification and precipitation to remove polymerase chain reaction (PCR) artifacts, samples were sequenced in multiplex on an Illumina HiSeq 2000. All target gene sequence processing was done with QIIME2 (Bolyen et al. 2019a, 2019b) via Qiita (Gonzalez et al. 2018). Raw sequencing data were trimmed and demultiplexed at 150 bases.

Amplicon sequence variants (ASV) were generated using the deblur algorithm (Amir et al. 2017). Phylogeny was created via SEPP (Janssen et al. 2018) within the QIIME2 fragment insertion plugin using default parameters. Taxonomy classification was done via the QIIME2 feature classifier plugin (Bokulich et al. 2018) and based on GreenGenes (McDonald et al. 2012) and Silva (Yilmaz et al. 2014). The resulting ASV table was filtered to remove any samples mislabeled with a probability above 0.20 using the sample type field as described by (Knights et al. 2011; Human Microbiome Project 2012). The resulting table was then rarefied at 10,000 sequences/sample to correct for uneven sequencing depth due to amplification differences between samples.

Alpha diversity is a within samples measure and was examined using evenness, observed OTU’s, and Faith’s Phylogenetic Diversity (Faith 1994). The UniFrac distance metric was used and is an algorithm that determines differences between microbial communities between samples (Beta-diversity) based upon their unique branch length on a phylogenetic tree (Lozupone and Knight 2005). Principal coordinate analysis (PCoA) using unweighted UniFrac distances (sensitive to rarer taxa) and weighted UniFrac distances (sensitive to abundances of taxa) are the best ways to visualize the microbiome as a whole. The prebiotic diet only altered unweighted UniFrac distances thus these PCoA graphs are depicted.

In addition to 16S rRNA gene sequencing, fecal samples collected on experimental day 42 were also analyzed using shotgun metagenomics. This approach allowed confirmation of the 16S rRNA gene findings with greater species level taxonomic resolution. Metagenomic sequence analysis was performed using Shogun with default parameters as described in (Hillmann et al. 2018). In short, adapter and host removal were performed using Atropos (Didion, Martin, and Collins 2017), then UTree was used to generate alignments against rep82, to finally generate taxonomic and pathway tables using the redistribute commands within Shogun.

### 2.10 Fecal Bile Acids

Fecal primary and bacterially modified secondary and conjugated bile acids were measured in all fecal samples using liquid chromatograph-tandom mass spectrometry (LC-MS/MS) and verified with bile acid standards as previously described in detail (Thompson et al. 2020). Sample ID’s were manually uploaded into an electronic spreadsheet and subsequently used to assign filenames during LC-MS/MS data acquisition. Fecal pellets were extracted as previously described (Melnik et al. 2017) and were weighed to 50.0 +/− 2 mg wet weight. The fecal homogenates were centrifuged at 14000 rpm for 15 min at 4°C. 1.2 mL aliquots were then transferred into Nunc 2.0 mL DeepWell plate (Thermo Catalog# 278743) and frozen at −80 °C prior to lyophilization using a FreeZone 4.5 L Benchtop Freeze Dryer with Centrivap Concentrator (Labconco). Wells were resuspended with 200 µL of resuspension solvent (50% MeOH spiked with 2.0 µM sulfadimethoxine), vortexed for 30 secs, and centrifuged at 2000 rpm for 15 min at 4 °C. 150 µL of the supernatant was transferred into a 96-well plate and maintained at 4 °C prior to LC-MS/MS analysis. A resuspension solvent QC and a six standard mix QC (50% MeOH spiked with 1.0 µM sulfamethazine, 1.0 µM sulfamethizole, 1.0 µM sulfachloropyridazine, 1.0 µM amitrypline, and 1.0 µM coumarin 314) was run every12^th^ sample to assess sample background, carry over, chromatography behavior, peak picking, and plate effects. Fecal extracts were analyzed using an ultra-high-performance liquid chromatography system (Vanquish, Thermo) coupled to a hybrid quadrupole-Orbitrap mass spectrometer (Q-Exactive, Thermo) fitted with a HESI probe. Reverse phase chromatographic separation was achieved using a Kinetex C18 1.7 µm, 100 Å, 50 x 2.1 mm column (Phenomenex) held at 40°C with a flow rate of 0.5 mL/min. 5.0 µL aliquots were injected per sample/QC. The mobile phase used was (A) 0.1% formic acid in water and (B) 0.1% formic acid in acetonitrile. The elution gradient was: 5% B for 1 min, increased to 100% B in the next 8 min, held at 100% B for two min, returned to 5.0% B in 0.5 min, equilibrated at 5.0% B for two min. Positive electrospray ionization parameters were: sheath gas flow rate of 52 (arb. units), aux gas flow rate of 14 (arb. units), sweep gas flow rate of 3 (arb. units), spray voltage of 3.5 kV, capillary temperature of 270 °C, S-Lens RF level of 50 (arb. units), and aux gas heater temperature of 435 °C. Negative electrospray ionization parameters were the following: sheath gas flow rate of 52 (arb. units), aux gas flow rate of 14 (arb. units), sweep gas flow rate of 3 (arb. units), spray voltage of 2.5 kV, capillary temperature of 270 °C, S-Lens RF level of 50 (arb. units), and aux gas heater temperature of 435 °C. All mass spectrometry data can be found in the online mass spectrometry repository Massive (http://massive.ucsd.edu) using the following accession numbers: MSV000082582 and MSV000080628.

#### 2.10.1 Bile Acid Standards

Primary, secondary, conjugated, and unconjugated bile acids were purchased (Cayman Chemical, 1180 East Ellsworth Road, Ann Arbor, MI, 48108, USA) and used for Level 1 bile acid identification in fecal samples. Standards were solubilized to a final concentration of 10 μM in 50% MeOH prior to LC-MS/MS injection.

#### 2.10.2 Rationale for Fecal Collection Time Course

Fecal samples collected on experimental days -33, 0, and 42 were analyzed. These samples were chosen based on the following rationale: Experimental day -33 (2 days on diet) established baseline microbiome and/or bile acid profiles; experimental day 0 (5 weeks on diet), and experimental day 42 (11 weeks on diet) revealed the stability of potential changes in the microbiome and bile acid profiles due to prebiotic diet consumption. Because our study included both prebiotic and control diet conditions, this time course enabled us to control for changes in the microbiome and bile acid profiles due to the rat age, surgery, and time in the vivarium. This is important because each of these factors can impact the fecal microbiome and bile acid profile (Yatsunenko et al. 2012; Perino et al. 2020; Keskey et al. 2020; Ferrie et al. 2020; Kaczmarek et al. 2018).

Fecal sample collections from both the CDR and NLD groups were matched for diurnal clock time. This is important because there is evidence that the microbiome and bile acid profiles differ across the diurnal cycle (Eggink et al. 2017; Govindarajan et al. 2016; Ovacik et al. 2010; Lundasen et al. 2006). The CDR group experienced repeated, weekly reversals of the light/dark cycle. On experimental day 42, the CDR group was mostly realigned to the undisturbed or normal light/dark cycle (see Figure 2H). The day 42 fecal samples, therefore, reveal if a *history* of CDR was sufficient to disturb the microbiome and/or bile acid profile without the confounding factor of diurnal rhythm differences (Tahara et al. 2018; Tognini et al. 2017; Leone et al. 2015).

**FIG. 2.**
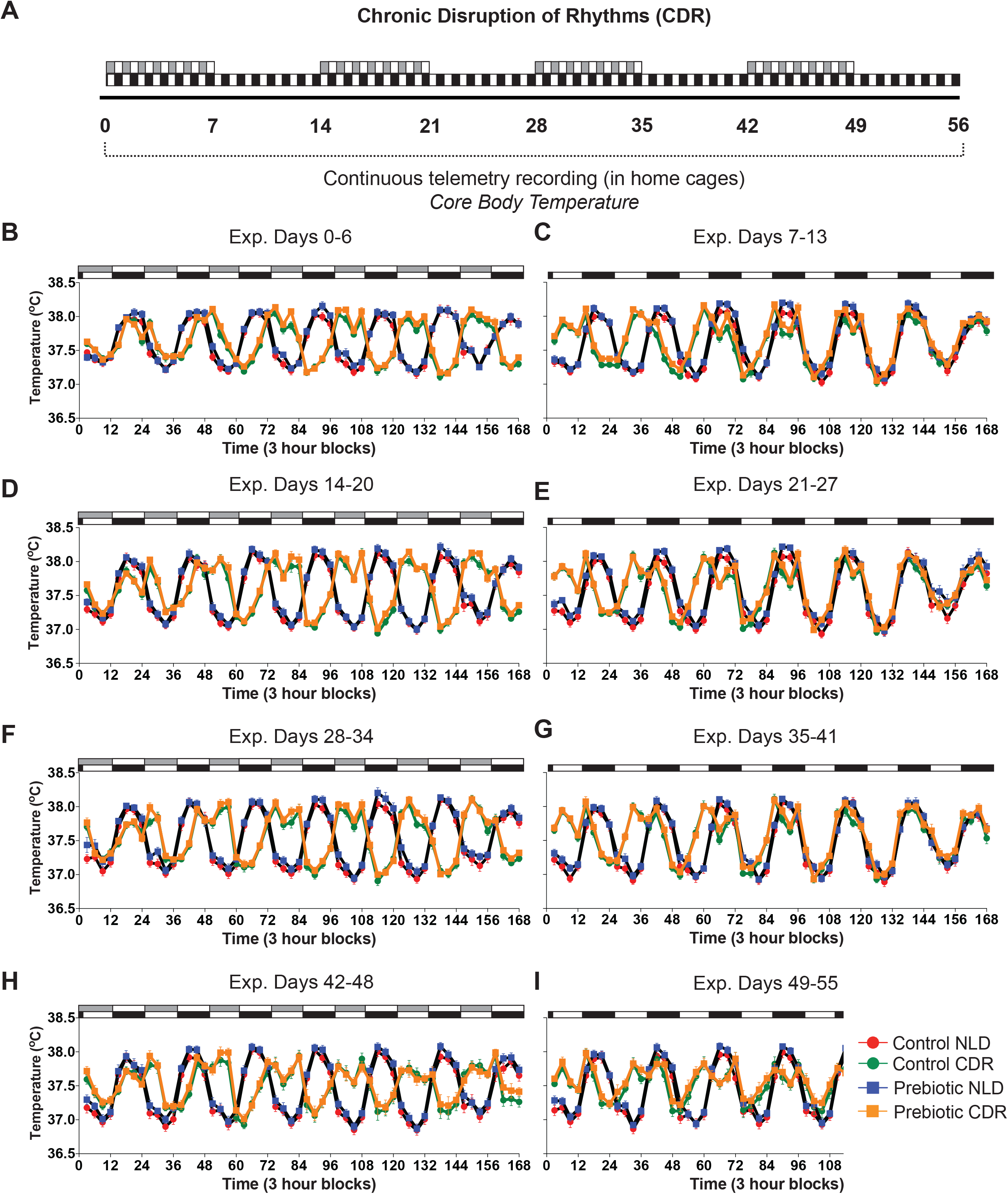
Core body temperature (CBT) data demonstrating forced misalignment of diurnal CBT across 56 experimental days due to weekly 12-hr light/dark reversals. On experimental days 0-6, 14-20, 28-43, and 42-48 the lights/dark cycle was reversed for the CDR group. Black and white bars represent the normal light/dark cycle (C, E, G, I), while the gray and white bars represent the reverse light dark cycle for rats exposed to CDR (B, D, F, H). (Control NLD n = 19; Control CDR n = 21; Prebiotic NLD n = 20; Prebiotic CDR n = 21).

### 2.11 Statistical Analysis

Data were analyzed using IBM SPSS statistics software version 26 and QIIME2. Data were examined using repeated measures ANOVA, one-way, and two-way ANOVA. When appropriate, post-hoc analysis for host physiological variables was performed using Fisher’s Least Significant Difference. In order to investigate differential abundance of species/genus level taxa between diet or CDR, an analysis of the composition of the microbiome (ANCOM) was performed (Mandal et al. 2015). For the gut microbiome analysis of Unifrac distance matrices, permutation multivariate analysis of variance (PERMANOVA) was used (Kelly et al. 2015; Tang, Chen, and Alekseyenko 2016). Stepwise multiple regression analyses were used to examine potential relationships between significant gut microbial changes, bile acids, and host physiological changes in dependent variable outcomes. The final group numbers for all dependent variables are presented in the Figure captions and ranged between 17 and 21 rats per group depending on experimental day and dependent measure. Group numbers varied due to the following: 1) given the length of the experiment several animals destroyed the EEG leads to the skull rendering the EEG/sleep data unusable; 2) fecal samples were excluded for not meeting quality control standards for microbiome sequencing (see microbiome methods); and 3) animals lacking complete data sets across all dependent measures were excluded from the regression analyses. Multiple correlations conducted on bile acid profiles and diet-impacted gut microbiota were corrected using Holm-Bonferroni. Two-tailed alpha was set at p < 0.05.

## 3. Results

### 3.1 Core Body Temperature (CBT) and Locomotor Activity (LA)

Depicted in Figure 2 are biotelemetry measures of core body temperature across the duration of the study (experimental days 0-55). Neither diet nor CDR impacted overall core body temperature. As expected after light/dark reversal, there were clear significant interactions of CDR across time on days 0-6, Figure 2B, (F_(55, 4235)_ = 190.429; p < 0.001); days 7-13m Figure 2C (F_(55, 4235)_ = 77.971; p < 0.001); days 14-20, Figure 2D (F_(55, 4235)_ = 251.456; p < 0.001); days 21-27, Figure 2E (F_(55, 4235)_ = 61.084; p = 0.001); on days 28-34, Figure 2F (F_(55, 4235)_ = 207.753; p < 0.001); days 35-41, Figure 2G (F_(55, 4235)_ = 66.984; p < 0.001); days 42-28, Figure 2H (F_(55, 4235)_ = 81.189; p < 0.001), and days 49-55, Figure 2i (F_(55, 4235)_ = 32.203; p < 0.001).

Depicted in Figure 3 are biotelemetry measures of locomotor activity across the duration of the study (experimental days 0-55). Neither diet nor CDR impacted overall locomotor activity (LA) on experimental days 0-6 and 28-34; however, in contrast to CBT, CDR increased overall LA across days 7-13, Figure 3C (F_(1,78)_ = 9.576; p = 0.002); days 14-20, Figure 3D (F_(1,78)_ = 4.133; p = 0.045); days 21-27, Figure 3E (F_(1,78)_ = 5.718; p = 0.019); days 35-41, Figure 3G (F_(1,78)_ = 13.104; p = 0.0005); days 42-48, Figure 3H (F_(1,78)_ = 14.62; p = 0.0003); and days 49-55, Figure 3i (F_(1,78)_ = 19.715; p < 0.0001). As expected after light/dark reversal, there were time by CDR interactions on experimental days 0-6, Figure 3B (F_(55, 4235)_ = 33.814 ; p < 0.001); days 7-13, Figure 3C (F_(55, 4235)_ = 45.299; p < 0.001); days 14-20, Figure 3D (F_(55, 4235)_ = 43.268; p < 0.001); days 21-27, Figure 3E (F_(55, 4235)_ = 34.756; p < 0.001); days 28-34, Figure 3F (F_(55, 4235)_ = 42.456; p < 0.001); days 35-41, Figure 3G (F_(55, 4235)_ = 30.256; p < 0.001); days 42-48, Figure 3H (F_(55, 4235)_ = 24.561; p < 0.001) and days 49-55, Figure 3i (F_(55, 4235)_ = 14.089; p < 0.001.

**FIG. 3.**
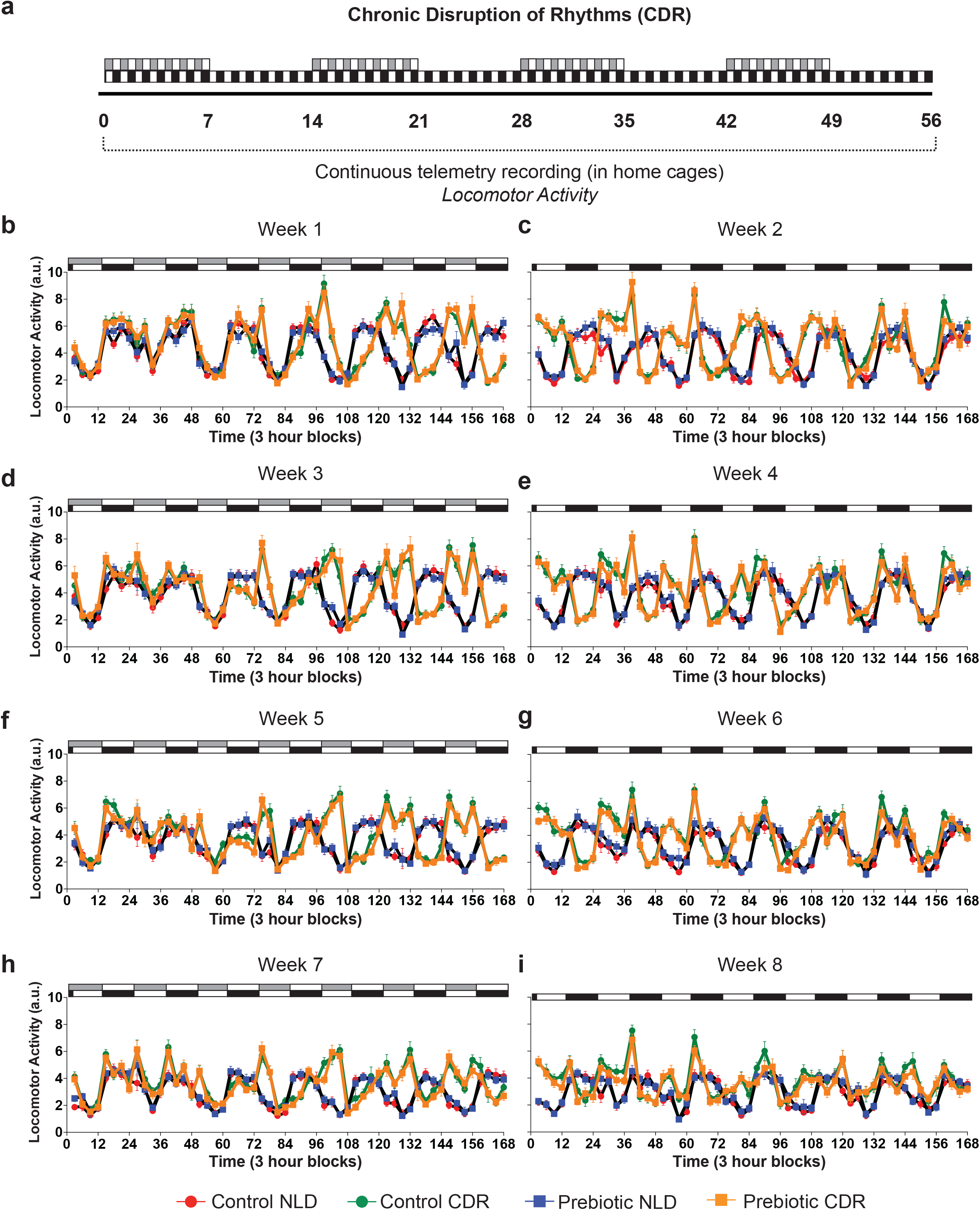
Locomotor Activity (LA) data demonstrating forced misalignment of diurnal LA across 56 experimental days and increased absolute LA in rats exposed to weekly 12-hr light/dark reversals (i.e., chronic disruption of rhythms (CDR)). On experimental days 0-6, 14-20, 28-43, and 42-48 the lights/dark cycle is reversed for the CDR group. Black and white bars represent the normal light/dark cycle (C, E, G, I), while the gray and white bars represent the reverse light/dark cycle for rats exposed to CDR (B, D, F, H). (Control NLD n = 19; Control CDR n = 21; Prebiotic NLD n = 20; Prebiotic CDR n = 21).

### 3.2 Prebiotic diet (GOS/PDX) alters the gut microbiome across time and increases the relative abundances of gut microbial species

16S rRNA gene microbiome analyses of fecal samples collected on experimental day -33, 0, and 42 were conducted using Analysis of the Composition of the Microbiome (ANCOM) (Mandal et al. 2015). As depicted in Figure 4, diet produced increases in the relative abundance of *Parabacteroides distasonis* on experimental days -33, Figure 4B (F_(1,37)_ = 2346.871; p < 0.001); day 0, Figure 4C (F_(1,73)_ = 121.058; p < 0.001); and day 42, Figure 4D (F_(1,78)_ = 235.977; p < 0.001). The relative abundance of *Ruminiclostridium 5* was also increased by diet on experimental day 0, Figure 4F (F_(1,73)_ = 101.416; p < 0.001) and day 42, Figure 4G (F_(1,78)_ = 154.75; p < 0.001); but not experimental day -33, Figure 4E. ANCOM did not reveal any differences in fecal bacteria collected from rats with a history of CDR compared to NLD.

**FIG. 4.**
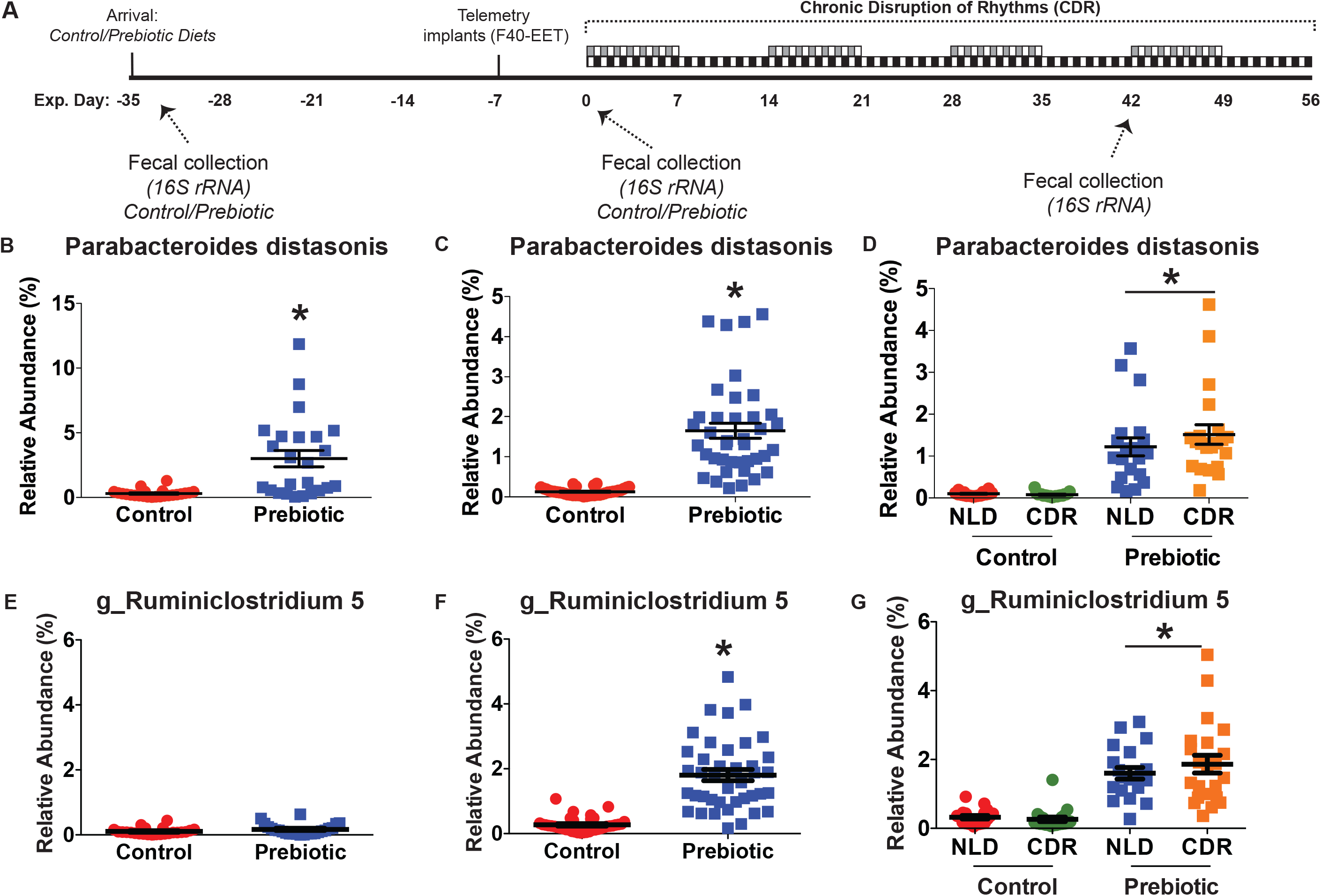
(A) Experimental timeline for fecal sample collections and 16S rRNA gene analyses. Prebiotic diet increased the relative abundance of *Parabacteroides distasonis* on (B) experimental day -33 that persisted in fecal samples collected on experimental days 0 (C) and 42 (D). The genus *Ruminiclostridium 5* was unchanged on experimental day -33 (E), and reliably elevated on experimental days 0 (F) and 42 (G). (Control NLD n = 20; Control CDR n = 21; Prebiotic NLD n = 20; Prebiotic CDR n = 21). Circle and square symbols represent individual rats for control and prebiotic diet respectively; horizontal line with error bars represent mean and standard error, * p < 0.05 compared to control diet where horizontal line under * represents significant main effect of diet.

Results from shotgun metagenomics analyses of experimental day 42 fecal samples are depicted in Figure 5. Principal coordinates analysis of the microbiome revealed a significant effect of prebiotic diet (PERMANOVA pseudo-F_(4,78)_ = 1.239; p = 0.00099) and not a history of CDR (Figure 5B), confirming the 16S rRNA gene results. The relative abundances of both *Parabacteroides distasonis* (F _(1,77)_ = 178.014; p < 0.0001; Figure 5F) and *Clostridium leptum* (F_(1,77)_ = 148.607; p < 0.0001; Figure 5G), a member of the *Ruminiclostridium* genera, were higher after prebiotic diet compared to control diet. These changes confirm our 16S rRNA gene sequencing results. Consistent with our 16S rRNA gene findings, ANCOM did not reveal any differences in fecal *Parabacteroides distasonis* and *Clostridium leptum* collected from rats with a history of CDR compared to NLD (Figure 5F, 5G).

**FIG. 5.**
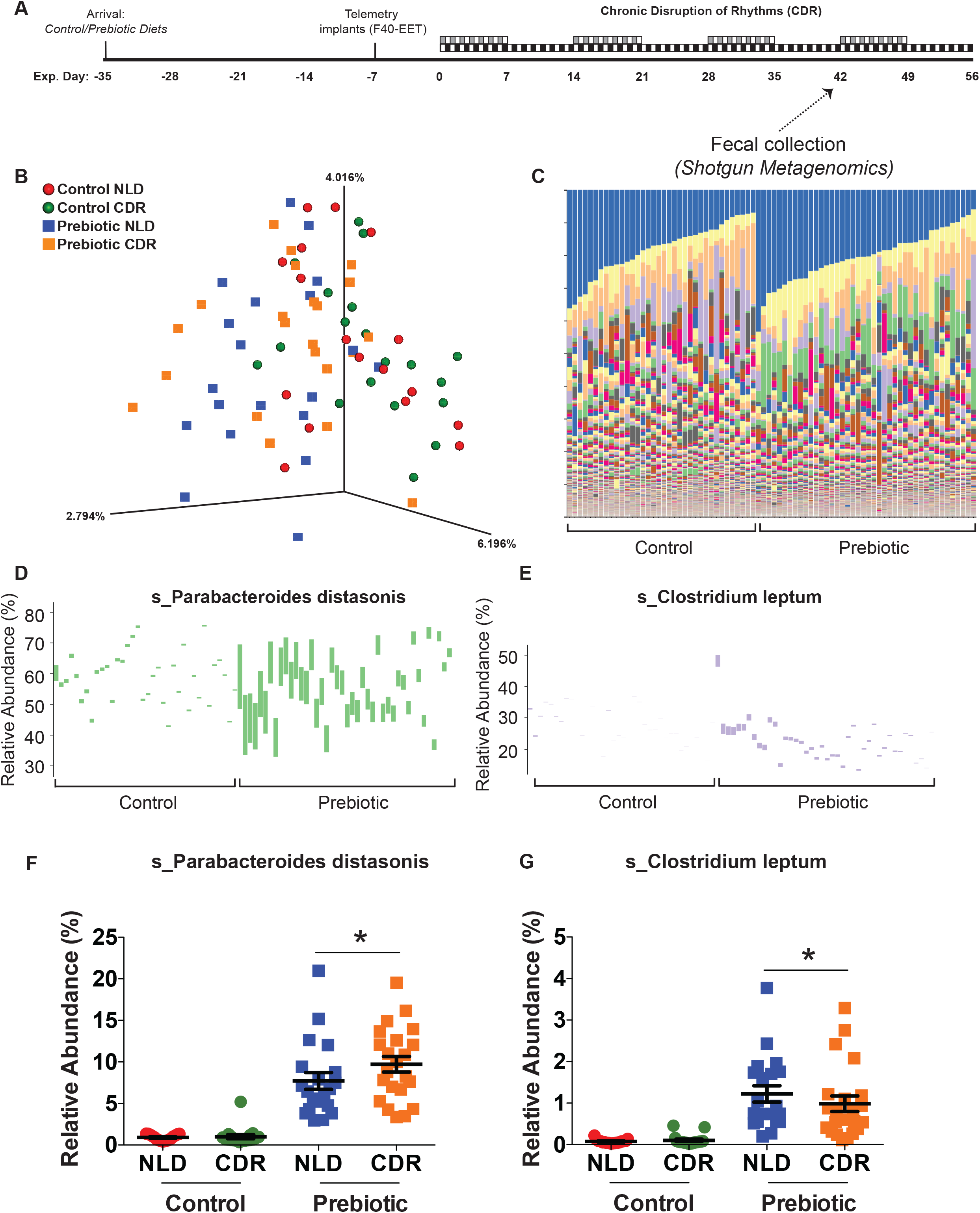
(A) Experimental timeline and shotgun metagenomics of fecal samples collected on experimental day 42. (B) Bacterial species level β-diversity PCoA capturing microbial ecology and (C) taxonomic classification where colors represent relative abundance of ∼4554 different (known/unknown) species. Vertical columns represent individual rats. The relative abundances of *Parabacteroides distasonis* (D, F) and *Clostridium leptum* (E, G) were higher in rats that consumed prebiotic diet. There was no effect of a history of CDR on these species. Horizontal line with error bars represents mean and standard error. ANCOM analysis, * p < 0.05 compared to control diet where horizontal line under * represents significant main effect of diet. (Control NLD n = 17; Control CDR n = 19; Prebiotic NLD n = 20; Prebiotic CDR n = 22).

### 3.3 Prebiotic diet (GOS/PDX) and a history of CDR alter bile acid profiles

All Level 1-identified bile acids measured in rat fecal samples collected on experimental days - 33, 0, and 42 are reported in Table 1. On experimental day -33, rats eating prebiotic diet compared to control chow had significantly lower beta muricholic acid (F_(1,46)_ = 6.537; p = 0.014), deoxycholic acid (F_(1,46)_ = 5.847; p = 0.02), glycoursodeoxycholic acid (F_(1,46)_ = 4.566; p = 0.038), and taurocholic acid (F_(1,46)_ = 8.988; p = 0.004). On experimental day 0, rats ingesting prebiotic compared to control diet had lower deoxycholic acid (F_(1,81)_ = 12.362; p = 0.001) and ursodeoxycholic acid (F_(1,81)_ = 8.61; p = 0.004) in fecal samples. On experimental day 42, prebiotic diet also reduced lithocholic acid (F_(3,81)_ = 6.265; p = 0.014) and ursodeoxycholic acid (F_(3,81)_ = 4.655; p = 0.034) while rats exposed to CDR compared to undisturbed NLD had higher levels of cholic acid (F _(3,81)_ = 7.54; p = 0.007) and taurocholic acid (F_(3,81)_ = 4.744; p = 0.032).

**Table 1.** Identified Fecal Bile Acids (numbers represent significant p-values; ns = not significant)

| Fecal Bile Acids |  |  |  |  |  |
| --- | --- | --- | --- | --- | --- |
| <u>Fecal Bile Acids</u> | <u>Diet (*p*-value)</u> |  |  | <u>CDR (*p*-value)</u> | <u>Effect</u> |
| Experimental Day | -33 | 0 | 42 | 42 |  |
| Primary Bile Acids |  |  |  |  |  |
| Alpha Muricholic Acid | 0.069 | ns | ns | ns | Trend Lower in Prebiotic |
| Beta Muricholic Acid | **0.014** | ns | ns | ns | Lower in Prebiotic |
| Cholic Acid | ns | ns | ns | **0.007** | Higher in CDR |
| Secondary Bile Acids |  |  |  |  |  |
| Deoxycholic Acid | **0.02** | **0.001** | ns | ns | Lower in Prebiotic |
| Hyodeoxycholic Acid | ns | ns | **0.054** | ns | Lower in Prebiotic |
| Lithocholic Acid | ns | ns | **0.014** | ns | Lower in Prebiotic |
| Ursodeoxycholic Acid | ns | **0.004** | **0.034** | ns | Lower in Prebiotic |
| Secondary Conjugated Bile Acids |  |  |  |  |  |
| Glycoursodeoxycholic Acid | **0.038** | ns | ns | ns | Lower in Prebiotic |
| Phenylalanine Conjugated Ursodeoxycholic Acid | ns | ns | ns | ns | No Effect |
| Tauroursodeoxycholic Acid | ns | ns | ns | ns | No Effect |
| Taurocholic Acid | **0.004** | ns | ns | **0.032** | Lower in Prebiotic / Higher in CDR |
| Glycocholic Acid | ns | ns | ns | ns | No Effect |

### 3.4 Significant associations between fecal bile acids and gut microbiota modulated by prebiotic diet (GOS/PDX)

The diagram in Figure 6 depicts significant relationships between *Parabacteroides distasonis, Ruminiclostridium 5*, primary, secondary, and conjugated bile acids in fecal samples collected on experimental day 0 (5 weeks on diet), prior to CDR exposure. *Parabacteroides distasonis* and *Ruminiclostridium 5* are positively correlated with each other and negatively correlated with taurocholic acid and deoxycholic acid, both microbially modified bile acids impacted by prebiotic diet.

**FIG. 6.**
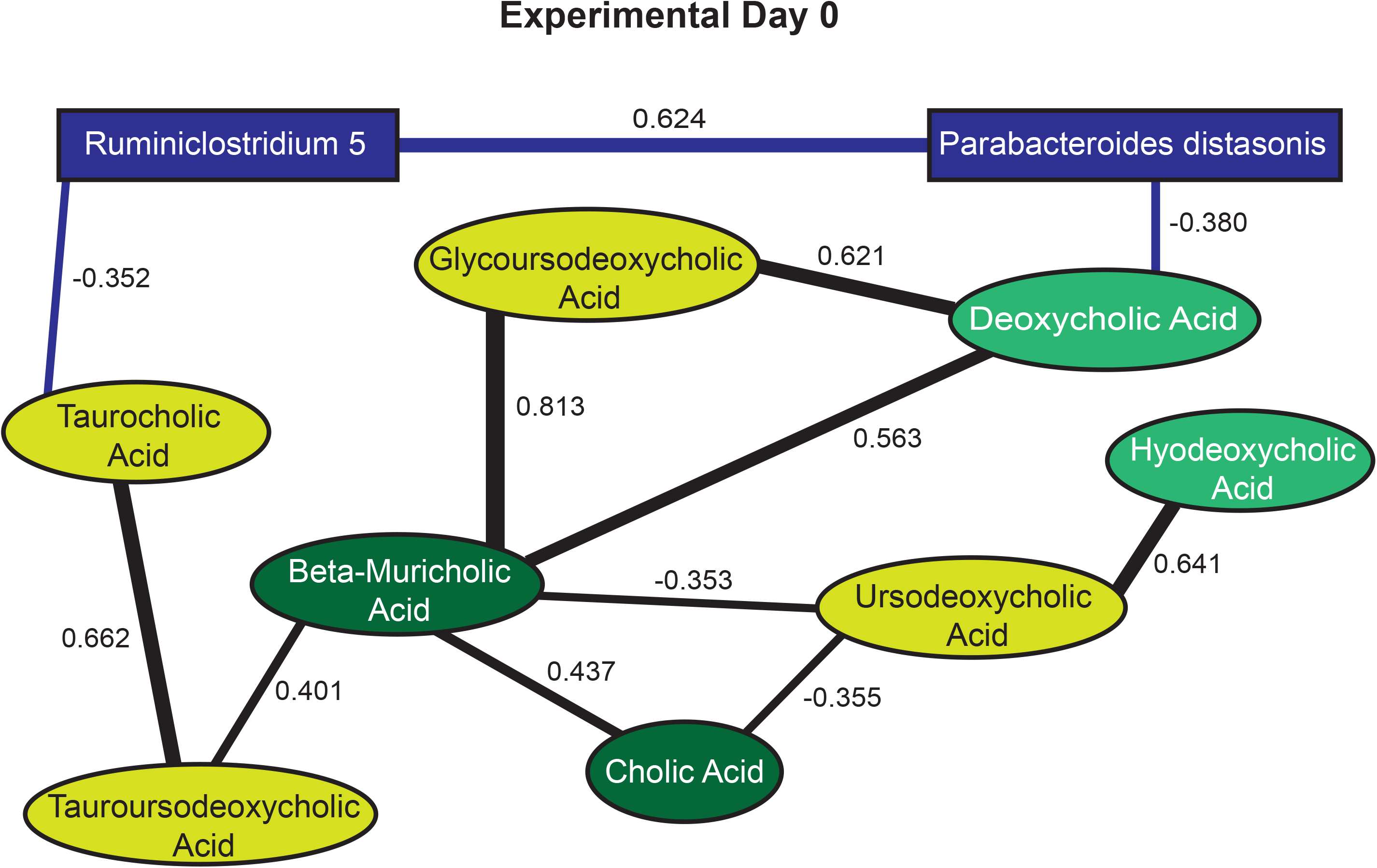
Diagram depicting significant relationships between microbes and bile acids in fecal samples collected on experimental day 0 from both prebiotic diet and control chow fed rats. Blue colored boxes represent gut bacteria, while dark green ovals represent primary bile acids, light green ovals represent secondary bile acids, and yellow ovals represent secondary conjugated bile acids. The lines connecting each box/circle represent the correlation between individual bacteria or bile acids. Pearson’s correlation coefficient (r) results were corrected using the Holm-Bonferonni method. (Control NLD n = 20; Control CDR n = 18; Prebiotic NLD n = 18; Prebiotic CDR n = 21).

To further elucidate the relationships between fecal bile acids, fecal microbes, and the potential impacts of diet and a history of CDR, stepwise regression analyses were performed (Figure 7). The following variables were included in the analyses: relative abundances of *Parabacteroides distasonis, Ruminiclostridium 5,* and bile acids in fecal samples collected on experimental day 0 and experimental day 42. Depicted in Figure 7, the relative abundances of *Parabacteroides distasonis* and bile acids reliably associate with the relative abundance of *Ruminiclostridium 5*.

**FIG. 7.**
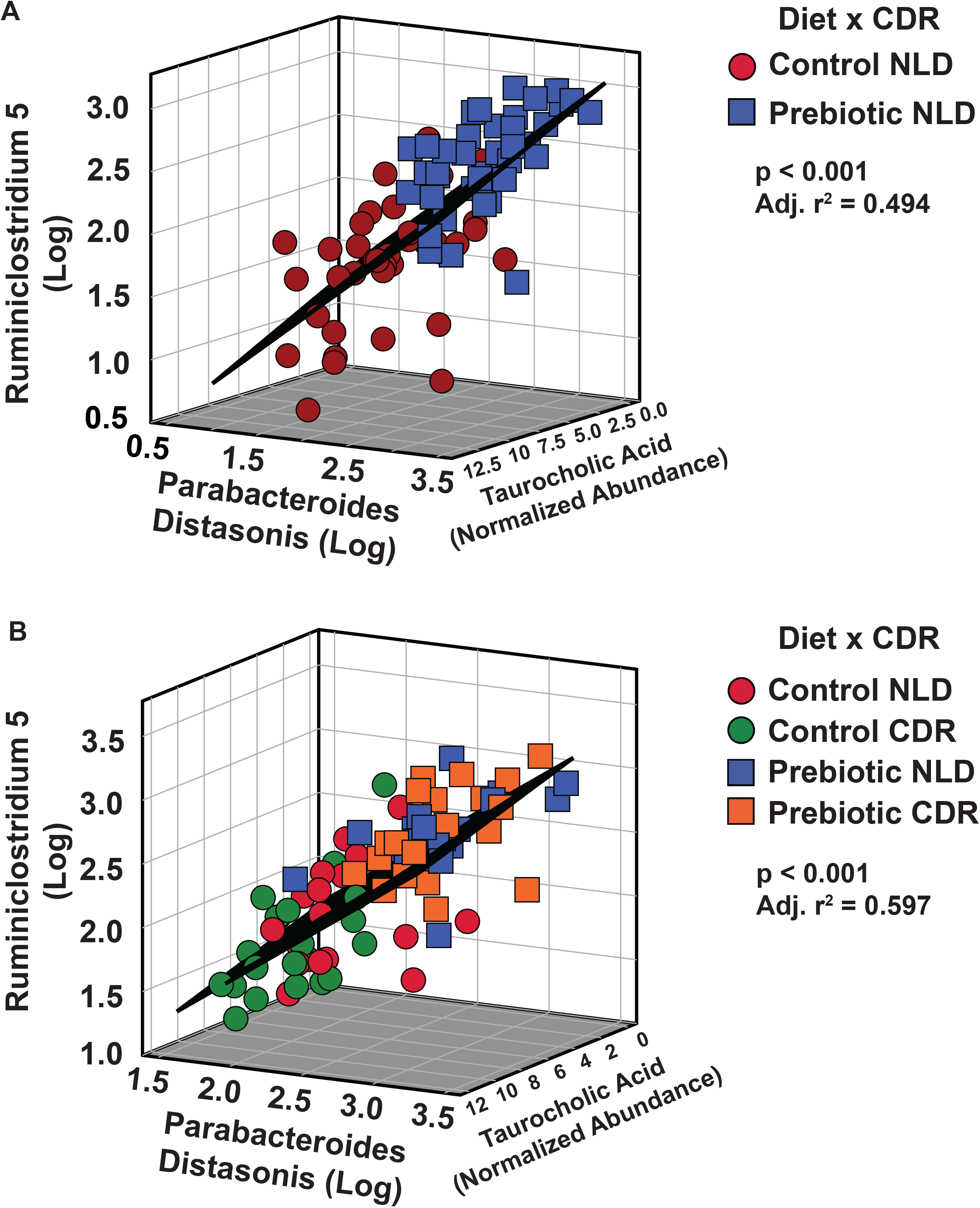
Regression analyses demonstrating that *Ruminiclostridium 5* is significantly associated with both *Parabacteroides distasonis* and taurocholic acid at both experimental day 0 (A), which is after 5 weeks on prebiotic diet and **prior to** CDR; and experimental day 42 (B) which is 11 weeks on prebiotic diet and **after** CDR. (Control NLD n = 20; Control CDR n = 18; Prebiotic NLD n = 18; Prebiotic CDR n = 21).

Figure 7A depicts the significant regression between *Ruminiclostridium 5*, *Parabacteroides distasonis,* and taurocholic acid (F_(2,75)_ = 37.674; p < 0.001; adj. r^2^ = 0.494). The regression equation is as follows: y = 0.603x_1_ – 0.051x_2_ + 0.992; where x_1_ = *Parabacteroides distasonis* and x_2_ = taurocholic acid. These relationships were also present in fecal samples collected on experimental day 42. Figure 7B depicts the significant regression between *Ruminiclostridium 5, Parabacteroides distasonis*, taurocholic acid and ursodeoxycholic acid (F _(3,80)_ = 40.568; p < 0.001; adj. r^2^ = 0.597; ursodeoxycholic acid not shown). The regression equation is as follows: y = 0.746x_1_ – 0.03x_2_ + 0.009x_3_; where x_1_ = *Parabacteroides distasonis* and x_2_ = taurocholic acid and x_3_ = ursodeoxycholic acid.

### 3.5 Prebiotic diet (GOS/PDX) and CDR impacts fecal microbiome alpha and beta diversity

Figure 8 depicts beta (unweighted UNIFRAC) and alpha diversity analyses of the fecal microbiome measured across time on diet and after CDR. Beta diversity was significantly different between prebiotic diet compared to control diet on experimental day 0 (pseudo-F_(2,78)_ = 4.69; p = 0.001; q = 0.001; Figure 8C) and experimental day 42 (pseudo-F_(4,83)_ = 2.077; p = 0.001; q = 0.0015; Figure 8D) but not experimental day -33 (Figure 8B). A history of CDR on experimental day 42 did not impact beta diversity based on unweighted or weighted UNIFRAC analyses; and diet did not impact beta diversity based on weighted UNIFRAC analysis at any time point measured. Importantly, findings based on 16S rRNA gene sequencing (Figure 8) and shotgun metagenomic sequencing (Figure 5) of fecal samples collected on experimental day 42 were consistent, i.e., diet, but not a history of CDR, significantly impacted beta diversity. Alpha diversity evenness was significantly different between prebiotic and control diet on experimental day 0 (F_(1,76)_ = 16.797; p = 0.0001; Figure 8F) and experimental day 42 (F_(1,81)_ = 5.006; p = 0.003; Figure 8G) but not experimental day -33 (Figure 8 E). On experimental day 42, a history of CDR reduced alpha diversity evenness in the control diet rats and prebiotic diet prevented that effect (F _(3,81)_ = 6.564; p = 0.012; see Figure 8G for results of posthoc test). There were no significant effects of diet or CDR on alpha diversity based on faith_PD or observed OTU’s analyses (data not shown).

**FIG. 8.**
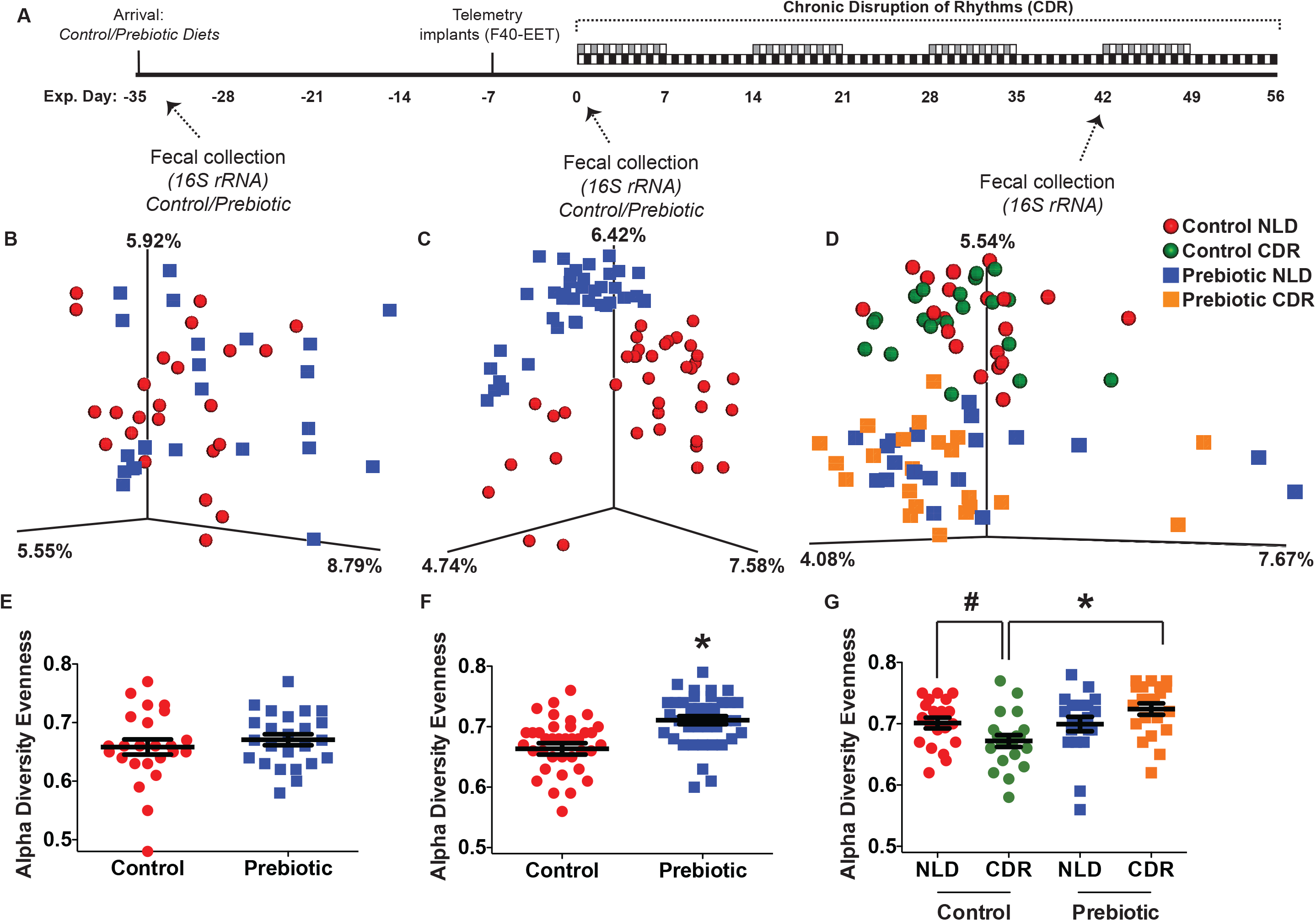
Unweighted Unifrac Principal coordinates analysis (PCoA) plots revealed that diet reliably altered β-diversity (B-D) and alpha diversity evenness (E-G) at experimental day 0 and 42, but not experimental day -33. G) A history of CDR reduced alpha diversity evenness in rats eating the control diet. This effect was prevented in rats eating prebiotic diet. * p < 0.05 compared to control (CDR) diet; # p < 0.05 compared to control diet NLD. (Control NLD n = 20; Control CDR n = 21; Prebiotic NLD n = 20; Prebiotic CDR n = 21).

### 3.6 Prebiotic diet (GOS/PDX) facilitates sleep recovery in rats with a history of CDR

Baseline sleep (%) measured on experimental day -1 was not impacted by diet (data not shown; NREM sleep F_(1, 71)_ = 0.08; p = 0.776; REM sleep F_(1, 71)_ = 0.07; p = 0.797; or Wake F_(1, 71)_ = 0.04; p = 0.837).

Diurnal sleep EEG data were analyzed for the first 36 hours after rats on the reverse light/dark cycle (CDR) were returned to the same light/dark cycle as the NLD rats on experimental days 8, 22, 36, and 50. Sleep architecture was characterized by the percentage of time spent in NREM (Figure 9A-D), REM (Figure 9E-H) and Wake (Figure 9I-L) depicted in 12-hr averages across the 36 hours. The number and duration of bouts in these sleep states are presented in Supplemental Figures 2 and 3, respectively.

**FIG. 9.**
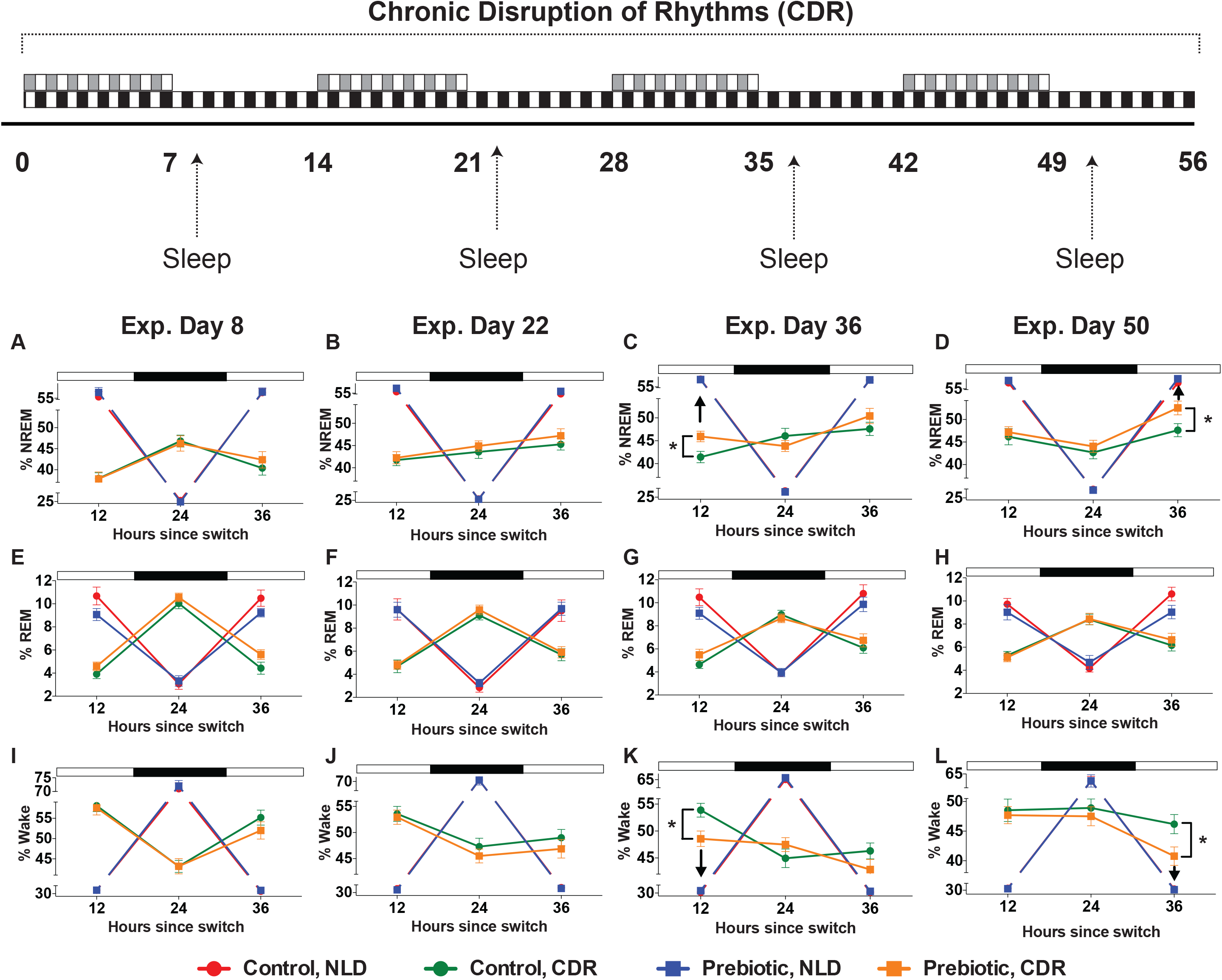
Sleep data (% time in sleep state) demonstrating that there were no diet effects on NREM on experimental days 8 (A) and 22 (B). Consumption of dietary prebiotics facilitated NREM sleep recovery following exposure to CDR. Denoted by the black arrows, rats eating prebiotic diet had greater NREM sleep recovery and spent less time awake in the light cycle on experimental days 36 (C, K) and 50 (D, L). * p < 0.05 compared to control diet CDR. (Control NLD n = 17; Control CDR n = 21; Prebiotic NLD n = 18; Prebiotic CDR n = 19).

Rats exposed to CDR had significant changes in sleep state, compared to NLD rats with undisturbed light/dark cycles (Figure 9). Consistent with the literature (Kervezee, Kosmadopoulos, and Boivin 2020), chronic disruption of diurnal rhythms reduces the percent time spent in NREM (Figures 9A-9D), and REM (Figures 9E-9H); and increases the percent time spent in Wake (Figures 9I-9L), for 36 hrs after returning to the original light/dark cycle on experimental days 8, 22, 36, and 50 (p < 0.05). There were also significant interactions between CDR, Diet, and hours (12h, 24h, 36h) for NREM and Wake, but not REM, on experimental days 36 and 50 (p < 0.05) but not days 8 and 22. Posthoc analyses revealed that rats exposed to CDR eating prebiotic diet, compared to control diet, spent more time in NREM sleep during the inactive/light cycle, 12 hrs since light/dark switch on experimental day 36 (Figure 9C; Fischer’s PLSD, p = 0.03) and 36 hrs since light/dark switch on experimental day 50 (Figure 9D; Fischer’s PLSD, p = 0.004). This same pattern occurred for the percent time in Wake. Rats exposed to CDR eating prebiotic diet compared to control diet, spent less time in Wake during the light/inactive cycle 12 hrs since light/dark switch on experimental days 36 (Figure 9K; Fischer’s PLSD, p = 0.04) and 36 hrs since light/dark switch on experimental day 50 (Figure 9L; p = 0.01), but not experimental days 8 and 22.

### 3.7 Prebiotic diet facilitates realignment of CBT bathyphase

To determine the impact of prebiotic diet on rate of diurnal realignment after chronic disruption of diurnal rhythms, we examined the circadian marker of bathyphase for CBT (Figure 10B) and LA (Figure 10C) across time and quantified the number of hours it took rats with a history of CDR to realign to their prior weeks’ bathyphase. Consumption of prebiotic diet compared to control diet resulted in a significant decrease in the time it took for the CBT bathyphase to realign in rats exposed to CDR (F_(1,38)_ = 5.941; p = 0.019; Figure 10D) and there was a significant decrease across time (F_(7,266)_ = 2.943; p < 0.0001; Figure 10D). When averaged, prebiotic compared to control diet, facilitated realignment by ∼17 hours (F_(1,38)_ = 5.941; p = 0.019; Figure 10F). Prebiotic did not reliably impact the rate of realignment for Locomotor Activity bathyphase (Figure 10E and 10G), but there was a significant decrease across time (F_(7,266)_ = 4.573; p < 0.001; Figure 10E).

**FIG. 10.**
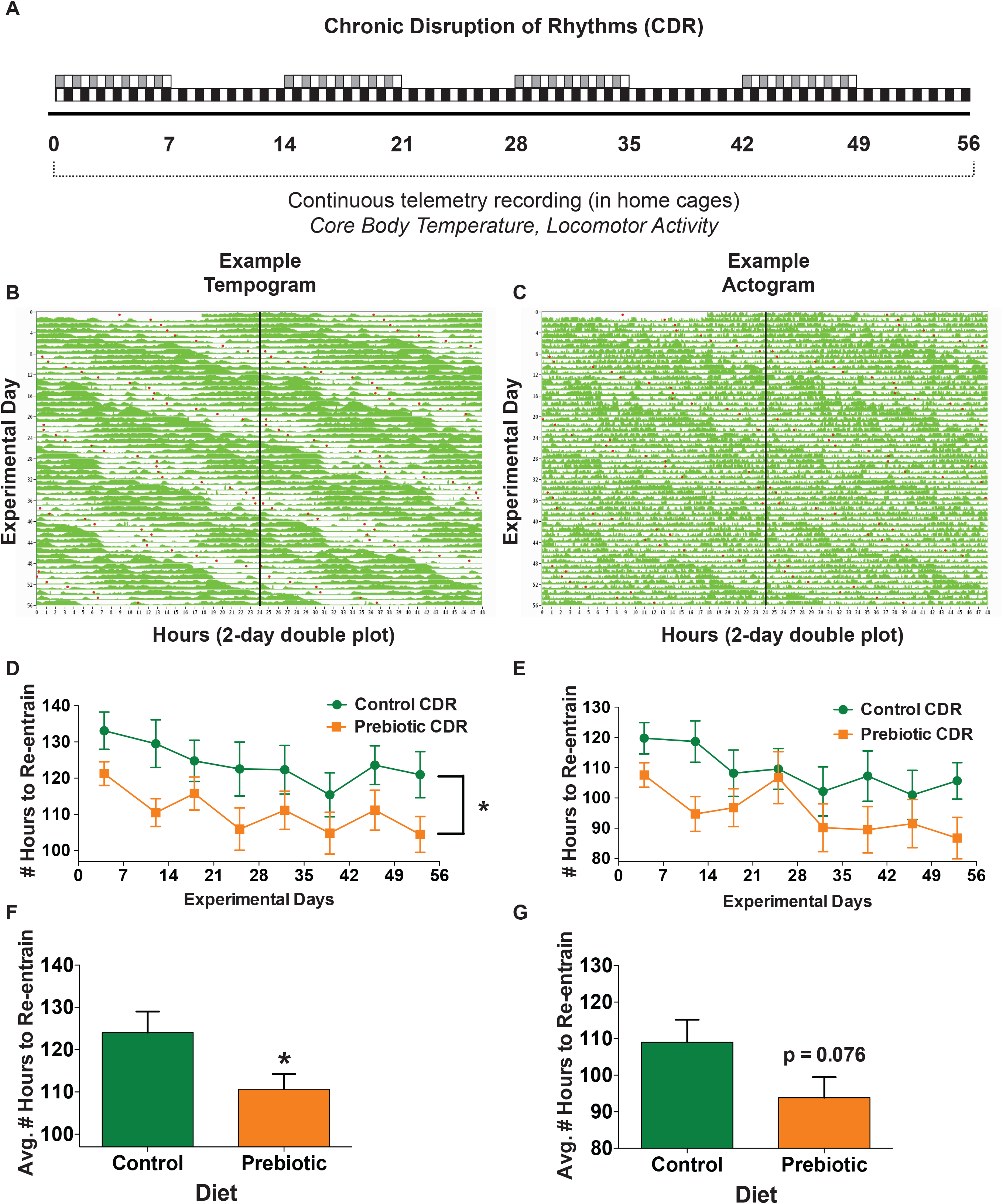
Core body temperature and locomotor activity bathyphase (diurnal minimum) data realignment abbreviated experimental timeline (A). Representative tempogram (B) and actogram (C). CBT/LA bathyphase automarked (red dots) using ClockLab software. CDR rats eating prebiotic compared to control diet more quickly realigned core body temperature (D, F), but not (E, G) locomotor activity bathyphase. (Control CDR n = 20; Prebiotic CDR n = 20).

### 3.8 CDR induced alterations to host physiology are related to *Ruminiclostridium 5* and fecal cholic/taurocholic acid

Based on our results describe above, rats consuming prebiotic diet compared to control diet, 1) more quickly realigned to core body temperature and sleep/wake rhythms to a new light/dark cycle; 2) increased the relative abundances of fecal *Parabacteroides distasonis* and *Ruminiclostridium 5* in the fecal microbiome; and 3) modulated fecal bile acid profiles. Using stepwise regression, therefore, we explored if the state of the fecal microbiome/bile acids *prior* to exposure to CDR would predict later impacts on time to realign CBT and facilitate NREM sleep recovery. Figure 11A depicts the significant regression where both cholic acid and *Ruminiclostridium 5* were related to CBT realignment (F_(2,36)_ = 7.376; p = 0.002; adj. r^2^ = 0.26). The regression equation is as follows: y = 5.562x_1_ – 12.832x_2_ + 77.922; where x_1_ = cholic acid and x_2_ = *Ruminiclostridium 5*.

**FIG. 11.**
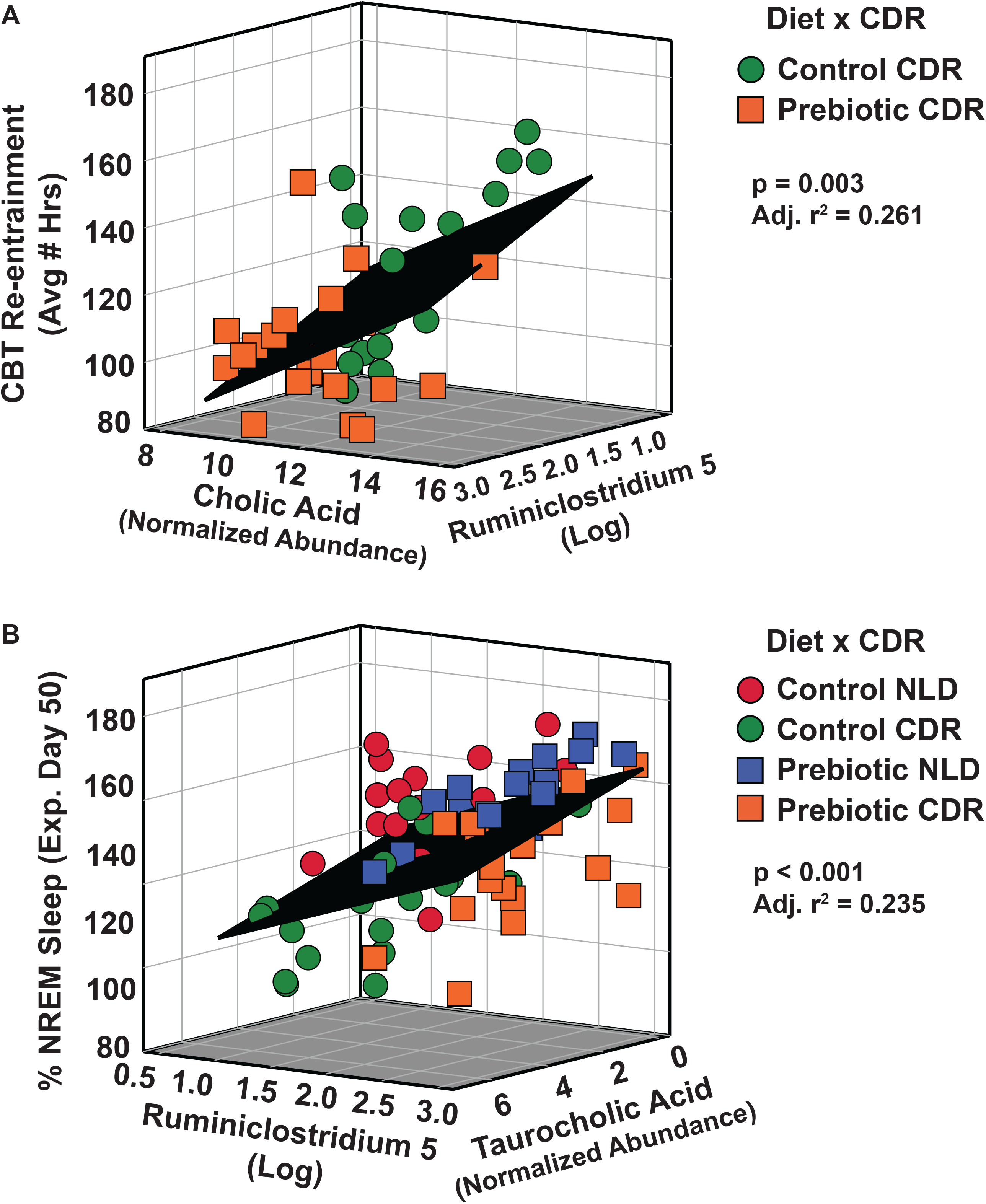
(A) Both cholic acid and *Ruminiclostridium 5* prior to CDR (experimental day 0) were related to time to realign CBT to the new light/dark cycle after CDR. (B) Both *Ruminiclostridium 5* and taurocholic acid prior to CDR (experimental day 0) were related to NREM sleep recovery after CDR (experimental day 50). (Control NLD n = 17; Control CDR n = 18; Prebiotic NLD n = 18; Prebiotic CDR n = 19).

Figure 11B depicts the significant regression where both *Ruminiclostridium 5* and taurocholic acid were related to facilitated NREM sleep recovery (F_(2,67)_ = 11.299; p < 0.001; adj. r^2^ = 0.235). The regression equation is as follows: y = 4.854x_1_ – 0.586x_2_ + 47.405; where x_1_ = *Ruminiclostridium 5* and x_2_ = taurocholic acid.

### 3.9 Body weights and food consumption

There were no differences in body weight (F _(1,82)_ = 0.014; p = 0.905; Control 140.8g ±3 8.5, Prebiotic 141.9g ± 45.7) or food consumption (F _(1,82)_ = 0.014; p = 0.905; Control 140.8g ± 38.5, Prebiotic 141.9g ± 45.7) the week before CDR exposure began. There were no effects of diet or CDR on body weight (Supplemental Figure 1B) or food consumption (Supplemental Figure 1C). All rats gained weight over the 8-weeks of CDR exposure (F_(7,560)_ = 1183.96; p < 0.0001; Supplemental Figure 1B).

## 4. Discussion

There is mounting evidence that diets enriched in GOS/PDX prebiotics modulate the fecal microbiome (Thompson et al. 2017; Mika et al. 2018), metabolome (Thompson et al. 2020), and stress physiology (Thompson et al. 2017; Mika et al. 2016). Based on 16S rRNA gene, shotgun metagenomics, LC-MS/MS with Level 1 bile acid identification, and *in vivo* biotelemetry analyses, we report that consumption of dietary prebiotics changes fecal microbial ecology, fecal bile acid profiles, and sleep/circadian physiology. Specifically, consumption of GOS/PDX: 1) rapidly and sustainably increases the relative abundances of *Parabacteroides distasonis* and genus *Ruminiclostridium 5*, of which the *Clostridium leptum* species belongs (Yutin and Galperin 2013); 2) reduces relative levels of fecal secondary bile acids, and 3) promotes sleep and CBT diurnal rhythm realignment after chronic disruption of rhythms.

Findings from prior research show that *Parabacteroides distasonis*, *Ruminiclostridium 5*, and *Clostridium leptum* are associated with positive health states, whereas reduced levels are associated with negative health states. Oral administration of *Parabacteroides distasonis,* for example, suppresses colon tumorigenesis and sustains epithelial barrier function in a mouse model (Koh et al. 2019). In addition, recovery from sleep fragmentation (Maki et al. 2020) is associated with increased relative abundance of *Ruminiclostridium 5*. In contrast, humans with metabolic syndrome had depleted relative abundances of *Parabacteroides distasonis*, which was reversed by a Mediterranean diet (Haro et al. 2016). Consumption of a high fat diet, considered to be a metabolic stressor, decreased the relative abundances of both *Parabacteroides distasonis* (Liu, Qin, and Wang 2019) and *Clostridium leptum* (Saha and Reimer 2014) in mice and rats, respectively. In addition, socially stressed mice had reduced levels of the genus Parabacteroides (Maltz et al. 2019) while psoriatic arthritis patients had decreased levels of *Parabacteroides distasonis* (Shapiro et al. 2019). Finally, a low relative abundance of *Ruminiclostridium 5* was reported in a rat model of acute pancreatitis (Chen, Huang, et al. 2017; Huang et al. 2017) and in patients with kidney stones (Tang et al. 2018). Our current results, taken together with previous findings, suggest that consumption of specific dietary prebiotics favors a health promoting gut microbial ecology.

Prebiotic diet also decreased the relative levels of several secondary and secondary conjugated bile acids and CDR significantly increased cholic/taurocholic acid in feces. The mechanisms responsible for changes in relative fecal bile acid levels are complex (Chiang 2013; Staley et al. 2017; Thomas et al. 2008). Lower secondary and secondary conjugated bile acids levels excreted in feces from rats consuming prebiotic diet compared to control diet, for example, could be due to changes in microbial deconjugation. We report compelling associations between *Ruminiclostridium 5*, *Parabacteroides distasonis* and taurocholic acid at multiple time points suggesting that reduced fecal levels of secondary/conjugated bile acids could be due to increases in microbial deconjugation and/or increases in reabsorption of conjugated bile.

Equally feasible, however, is that lower fecal secondary and secondary conjugated bile acids levels are due to faster and more efficient gut reabsorption. There is support for this idea in the literature. Bile acids are reabsorbed in the gut faster at lower pH (pH 7.0) versus higher pH (pH 8.0) and consumption of dietary prebiotic oligosaccharides, compared to a control diet, lowers the colonic pH from pH 8.0 to pH 7.0 (van Meer et al. 2008; Mekhjian, Phillips, and Hofmann 1979), which would result in faster intestinal reabsorption of conjugated bile acids and reduced fecal excretion.

Here we report that CDR increases relative levels of cholic acid and taurocholic acid in feces. There is previous evidence that CDR alters bile acid metabolism (Ho 1976; Yu et al. 2020; Ma et al. 2009). Ferrell and Chiang, for example, reported that 1-week of CDR changed bile acid composition in the intestines of female mice (Ferrell and Chiang 2015). In addition to CDR, rats exposed to chronic stress also had higher fecal cholic acid compared to control rats (Yu et al. 2017; Zheng et al. 2011). Higher cholic acid levels in colonic fecal matter could be especially problematic because cholic acid supports the germination of pathogenic *C. difficile* spores (Wilson 1983).

Reduced bile acid absorption, faster intestinal transit time, increases in *de novo* bile acid synthesis, or changes in FXR signaling (Silvennoinen et al. 2015) are all feasible mechanisms for increasing fecal levels of primary bile acids. A recent study by Silvennoinen et al., determined that fecal bile acid concentrations increased after chronic restraint stress were likely due to reduced bile acid absorption and faster intestinal transit time rather than changes in *de novo* bile acid synthesis or FXR expression (Silvennoinen et al. 2015). Our current results are consistent with previous findings and suggest that increased fecal output of bile acids, like cholic acid, may be indicative of a chronic stress-like state within the gut microbial-bile acid environment that could contribute to negative health outcomes.

## 4.1 Conclusions

A strength of the current study was that the fecal microbiome, fecal bile acids, CBT, and sleep were collected at several time points in the same rats. This greatly increases the taxonomic and temporal resolution of these measures and reveals meaningful associations between the microbiome, the metabolome, and sleep/circadian physiology. We report here that *Ruminiclostridium 5* and cholic/taurocholic acid *prior to* exposure to CDR were associated with facilitated realignment of core body temperature and NREM sleep recovery following the stress of CDR. These data suggest, therefore, that consumption of gut microbial modulating substrates, like prebiotics, may be a feasible way of promoting a stress robust phenotype.

---

**SUPPLEMENTAL FIG. 1.**
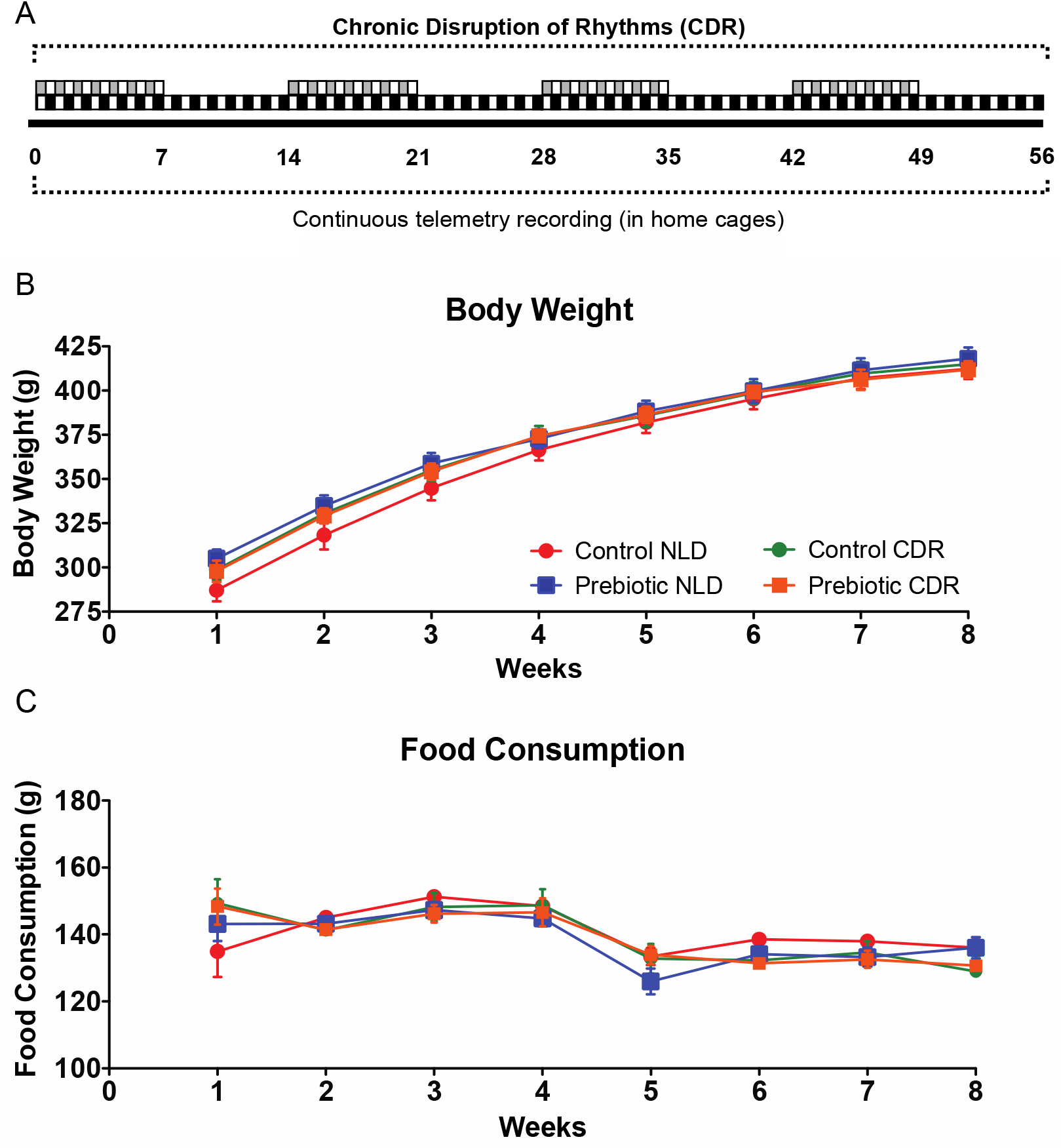
Body/Food weights. Abbreviated experimental timeline (A). Body weight (B) and food consumption (C) across 56 experimental days (8 weeks). All rats gained weight and neither diet nor chronic disruption of rhythms (CDR) affected body weight gain. Abbreviations: grams (g); normal light/dark cycle (NLD). #p < 0.05 vs. CDR. (Control NLD n = 20; Control CDR n = 21; Prebiotic NLD n = 20; Prebiotic CDR n = 21).

**SUPPLEMENTAL FIG. 2.**
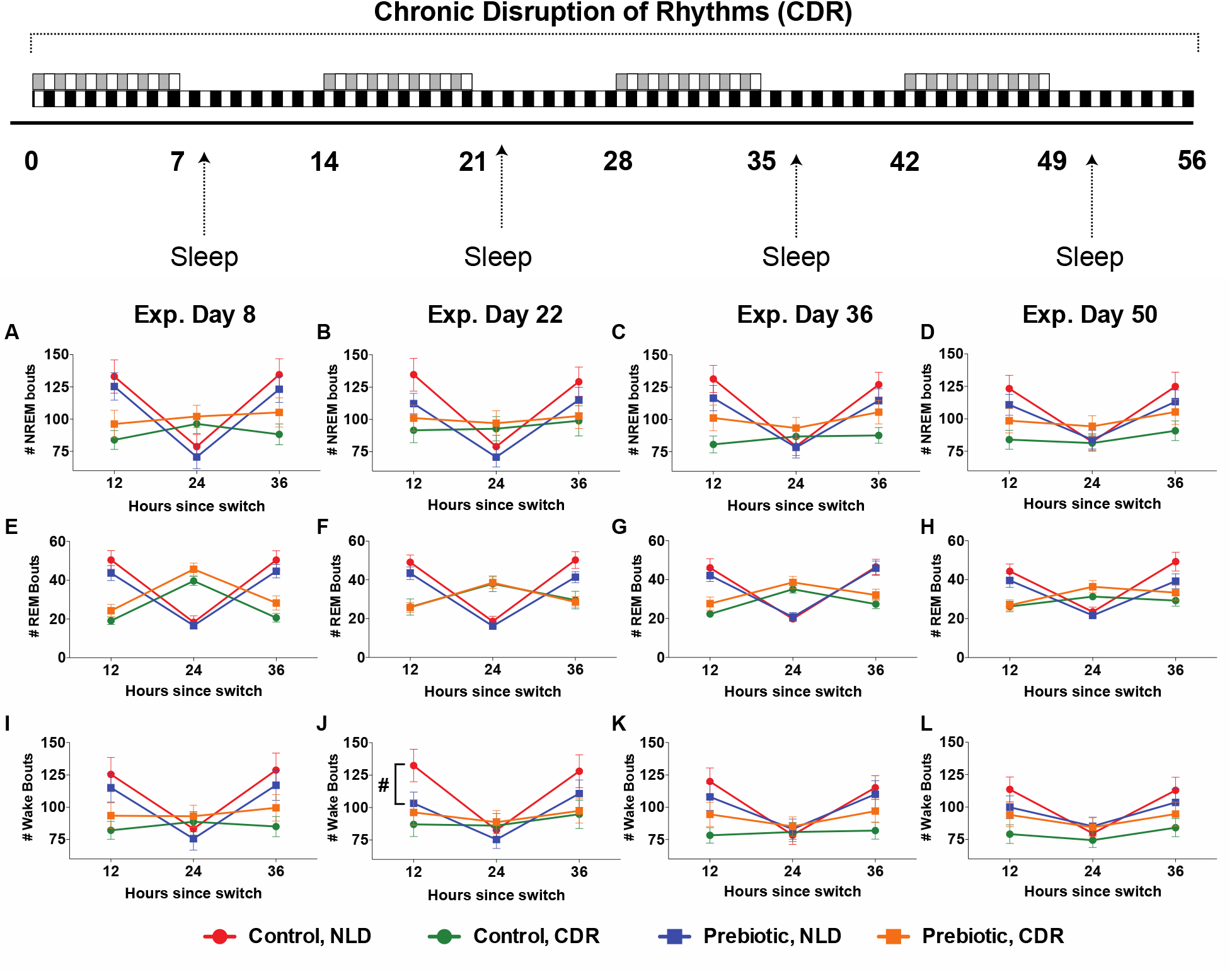
Diet had little impact on the number of NREM (A-D), REM (E-H) and wake (I-L) bouts after CDR. Rats housed in the NLD condition and eating prebiotic diet, however, had fewer wake bouts compared to control diet on experimental day 22 (J). (Control NLD n = 17; Control CDR n = 21; Prebiotic NLD n = 18; Prebiotic CDR n = 19).

**SUPPLEMENTAL FIG. 3.**
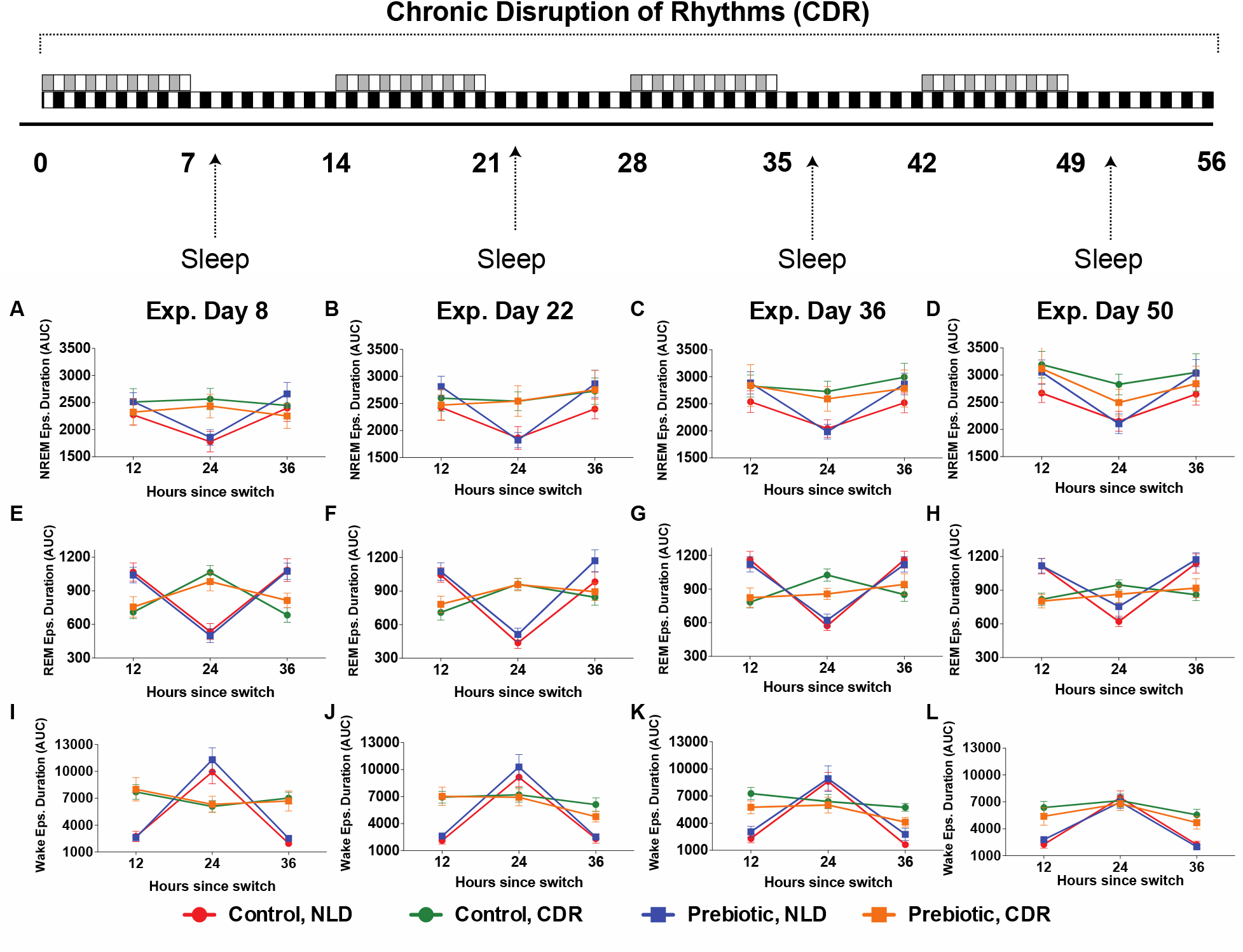
There were no significant effects of diet on NREM (A-D), REM (E-H) and wake (I-L) bout durations. (Control NLD n = 17; Control CDR n = 21; Prebiotic NLD n = 18; Prebiotic CDR n = 19).

## References

Al-Aqil, F. A., M. J. Monte, A. Peleteiro-Vigil, O. Briz, R. Rosales, R. Gonzalez, C. J. Aranda, B. Ocon, I. Uriarte, F. S. de Medina, O. Martinez-Augustin, M. A. Avila, J. J. G. Marin, and M. R. Romero. 2018. ‘Interaction of glucocorticoids with FXR/FGF19/FGF21-mediated ileum-liver crosstalk’, Biochim Biophys Acta Mol Basis Dis, 1864: 2927–37.

Amir, A., D. McDonald, J. A. Navas-Molina, E. Kopylova, J. T. Morton, Z. Zech Xu, E. P. Kightley, L. R. Thompson, E. R. Hyde, A. Gonzalez, and R. Knight. 2017. ‘Deblur Rapidly Resolves Single-Nucleotide Community Sequence Patterns’, mSystems, 2.

An, J., J. Zhao, and S. Liu. 2019. ‘Bone marrow-derived interstitial cells of cajal are increased by electroacupuncture in the colon of diabetic mice’, Journal of gastroenterology and hepatology.

Belfry, G. R., G. H. Raymer, G. D. Marsh, D. H. Paterson, R. T. Thompson, and S. G. Thomas. 2012. ‘Muscle metabolic status and acid-base balance during 10-s work:5-s recovery intermittent and continuous exercise’, J Appl Physiol (1985), 113: 410–7.

Benloucif, S., H. J. Burgess, E. B. Klerman, A. J. Lewy, B. Middleton, P. J. Murphy, B. L. Parry, and V. L. Revell. 2008. ‘Measuring melatonin in humans’, J Clin Sleep Med, 4: 66–9.

Bokulich, N. A., B. D. Kaehler, J. R. Rideout, M. Dillon, E. Bolyen, R. Knight, G. A. Huttley, and J. Gregory Caporaso. 2018. ’Optimizing taxonomic classification of marker-gene amplicon sequences with QIIME 2’s q2-feature-classifier plugin’, Microbiome, 6: 90.

Bolyen, E., J. R. Rideout, M. R. Dillon, N. A. Bokulich, C. C. Abnet, G. A. Al-Ghalith, H. Alexander, E. J. Alm, M. Arumugam, F. Asnicar, Y. Bai, J. E. Bisanz, K. Bittinger, A. Brejnrod, C. J. Brislawn, C. T. Brown, B. J. Callahan, A. M. Caraballo-Rodriguez, J. Chase, E. K. Cope, R. Da Silva, C. Diener, P. C. Dorrestein, G. M. Douglas, D. M. Durall, C. Duvallet, C. F. Edwardson, M. Ernst, M. Estaki, J. Fouquier, J. M. Gauglitz, S. M. Gibbons, D. L. Gibson, A. Gonzalez, K. Gorlick, J. Guo, B. Hillmann, S. Holmes, H. Holste, C. Huttenhower, G. A. Huttley, S. Janssen, A. K. Jarmusch, L. Jiang, B. D. Kaehler, K. B. Kang, C. R. Keefe, P. Keim, S. T. Kelley, D. Knights, I. Koester, T. Kosciolek, J. Kreps, M. G. I. Langille, J. Lee, R. Ley, Y. X. Liu, E. Loftfield, C. Lozupone, M. Maher, C. Marotz, B. D. Martin, D. McDonald, L. J. McIver, A. V. Melnik, J. L. Metcalf, S. C. Morgan, J. T. Morton, A. T. Naimey, J. A. Navas-Molina, L. F. Nothias, S. B. Orchanian, T. Pearson, S. L. Peoples, D. Petras, M. L. Preuss, E. Pruesse, L. B. Rasmussen, A. Rivers, M. S. Robeson, 2nd, P. Rosenthal, N. Segata, M. Shaffer, A. Shiffer, R. Sinha, S. J. Song, J. R. Spear, A. D. Swafford, L. R. Thompson, P. J. Torres, P. Trinh, A. Tripathi, P. J. Turnbaugh, S. Ul-Hasan, J. J. J. van der Hooft, F. Vargas, Y. Vazquez-Baeza, E. Vogtmann, M. von Hippel, W. Walters, Y. Wan, M. Wang, J. Warren, K. C. Weber, C. H. D. Williamson, A. D. Willis, Z. Z. Xu, J. R. Zaneveld, Y. Zhang, Q. Zhu, R. Knight, and J. G. Caporaso. 2019a. ‘Author Correction: Reproducible, interactive, scalable and extensible microbiome data science using QIIME 2’, Nat Biotechnol, 37: 1091.

Bolyen, E., J. R. Rideout, M. R. Dillon, N. A. Bokulich, C. C. Abnet, G. A. Al-Ghalith, H. Alexander, E. J. Alm, M. Arumugam, F. Asnicar, Y. Bai, J. E. Bisanz, K. Bittinger, A. Brejnrod, C. J. Brislawn, C. T. Brown, B. J. Callahan, A. M. Caraballo-Rodriguez, J. Chase, E. K. Cope, R. Da Silva, C. Diener, P. C. Dorrestein, G. M. Douglas, D. M. Durall, C. Duvallet, C. F. Edwardson, M. Ernst, M. Estaki, J. Fouquier, J. M. Gauglitz, S. M. Gibbons, D. L. Gibson, A. Gonzalez, K. Gorlick, J. Guo, B. Hillmann, S. Holmes, H. Holste, C. Huttenhower, G. A. Huttley, S. Janssen, A. K. Jarmusch, L. Jiang, B. D. Kaehler, K. B. Kang, C. R. Keefe, P. Keim, S. T. Kelley, D. Knights, I. Koester, T. Kosciolek, J. Kreps, M. G. I. Langille, J. Lee, R. Ley, Y. X. Liu, E. Loftfield, C. Lozupone, M. Maher, C. Marotz, B. D. Martin, D. McDonald, L. J. McIver, A. V. Melnik, J. L. Metcalf, S. C. Morgan, J. T. Morton, A. T. Naimey, J. A. Navas-Molina, L. F. Nothias, S. B. Orchanian, T. Pearson, S. L. Peoples, D. Petras, M. L. Preuss, E. Pruesse, L. B. Rasmussen, A. Rivers, M. S. Robeson, 2nd, P. Rosenthal, N. Segata, M. Shaffer, A. Shiffer, R. Sinha, S. J. Song, J. R. Spear, A. D. Swafford, L. R. Thompson, P. J. Torres, P. Trinh, A. Tripathi, P. J. Turnbaugh, S. Ul-Hasan, J. J. J. van der Hooft, F. Vargas, Y. Vazquez-Baeza, E. Vogtmann, M. von Hippel, W. Walters, Y. Wan, M. Wang, J. Warren, K. C. Weber, C. H. D. Williamson, A. D. Willis, Z. Z. Xu, J. R. Zaneveld, Y. Zhang, Q. Zhu, R. Knight, and J. G. Caporaso. 2019b. ‘Reproducible, interactive, scalable and extensible microbiome data science using QIIME 2’, Nat Biotechnol, 37: 852–57.

Brownawell, A. M., W. Caers, G. R. Gibson, C. W. Kendall, K. D. Lewis, Y. Ringel, and J. L. Slavin. 2012. ‘Prebiotics and the health benefits of fiber: current regulatory status, future research, and goals’, The Journal of nutrition, 142: 962–74.

Cai, Y., W. Liu, Y. Lin, S. Zhang, B. Zou, D. Xiao, L. Lin, Y. Zhong, H. Zheng, Q. Liao, and Z. Xie. 2018. ‘Compound polysaccharides ameliorate experimental colitis by modulating gut microbiota composition and function’, Journal of gastroenterology and hepatology.

Cardelle-Cobas, A., N. Corzo, A. Olano, C. Pelaez, T. Requena, and M. Avila. 2011. ‘Galactooligosaccharides derived from lactose and lactulose: influence of structure on Lactobacillus, Streptococcus and Bifidobacterium growth’, International journal of food microbiology, 149: 81–7.

Chang, M., Y. Zhao, G. Qin, and X. Zhang. 2018. ‘Fructo-Oligosaccharide Alleviates Soybean-Induced Anaphylaxis in Piglets by Modulating Gut Microbes’, Front Microbiol, 9: 2769.

Chen, D., X. Yang, J. Yang, G. Lai, T. Yong, X. Tang, O. Shuai, G. Zhou, Y. Xie, and Q. Wu. 2017. ‘Prebiotic Effect of Fructooligosaccharides from Morinda officinalis on Alzheimer’s Disease in Rodent Models by Targeting the Microbiota-Gut-Brain Axis’, Front Aging Neurosci, 9: 403.

Chen, J., C. Huang, J. Wang, H. Zhou, Y. Lu, L. Lou, J. Zheng, L. Tian, X. Wang, Z. Cao, and Y. Zeng. 2017. ‘Dysbiosis of intestinal microbiota and decrease in paneth cell antimicrobial peptide level during acute necrotizing pancreatitis in rats’, PloS one, 12: e0176583.

Chiang, J. Y. 2013. ‘Bile acid metabolism and signaling’, Compr Physiol, 3: 1191–212.

Chiang, J. Y., P. Pathak, H. Liu, A. Donepudi, J. Ferrell, and S. Boehme. 2017. ‘Intestinal Farnesoid X Receptor and Takeda G Protein Couple Receptor 5 Signaling in Metabolic Regulation’, Dig Dis, 35: 241–45.

Depner, C. M., E. R. Stothard, and K. P. Wright, Jr. 2014. ‘Metabolic consequences of sleep and circadian disorders’, Curr Diab Rep, 14: 507.

Didion, J. P., M. Martin, and F. S. Collins. 2017. ‘Atropos: specific, sensitive, and speedy trimming of sequencing reads’, PeerJ, 5: e3720.

Eggink, H. M., J. E. Oosterman, P. de Goede, E. M. de Vries, E. Foppen, M. Koehorst, A. K. Groen, A. Boelen, J. A. Romijn, S. E. la Fleur, M. R. Soeters, and A. Kalsbeek. 2017. ‘Complex interaction between circadian rhythm and diet on bile acid homeostasis in male rats’, Chronobiology international, 34: 1339–53.

Faith, D. P. 1994. ‘Phylogenetic pattern and the quantification of organismal biodiversity’, Philosophical transactions of the Royal Society of London. Series B, Biological sciences, 345: 45–58.

Ferrario, C., R. Statello, L. Carnevali, L. Mancabelli, C. Milani, M. Mangifesta, S. Duranti, G. A. Lugli, B. Jimenez, S. Lodge, A. Viappiani, G. Alessandri, M. Dall’Asta, D. Del Rio, A. Sgoifo, D. van Sinderen, M. Ventura, and F. Turroni. 2017. ‘How to Feed the Mammalian Gut Microbiota: Bacterial and Metabolic Modulation by Dietary Fibers’, Front Microbiol, 8: 1749.

Ferrell, J. M., and J. Y. Chiang. 2015. ‘Short-term circadian disruption impairs bile acid and lipid homeostasis in mice’, Cell Mol Gastroenterol Hepatol, 1: 664–77.

Ferrie, S., A. Webster, B. Wu, C. Tan, and S. Carey. 2020. ‘Gastrointestinal surgery and the gut microbiome: a systematic literature review’, European journal of clinical nutrition.

Gibson, G. R., R. Hutkins, M. E. Sanders, S. L. Prescott, R. A. Reimer, S. J. Salminen, K. Scott, C. Stanton, K. S. Swanson, P. D. Cani, K. Verbeke, and G. Reid. 2017. ‘Expert consensus document: The International Scientific Association for Probiotics and Prebiotics (ISAPP) consensus statement on the definition and scope of prebiotics’, Nat Rev Gastroenterol Hepatol, 14: 491–502.

Gonzalez, A., J. A. Navas-Molina, T. Kosciolek, D. McDonald, Y. Vazquez-Baeza, G. Ackermann, J. DeReus, S. Janssen, A. D. Swafford, S. B. Orchanian, J. G. Sanders, J. Shorenstein, H. Holste, S. Petrus, A. Robbins-Pianka, C. J. Brislawn, M. Wang, J. R. Rideout, E. Bolyen, M. Dillon, J. G. Caporaso, P. C. Dorrestein, and R. Knight. 2018. ‘Qiita: rapid, web-enabled microbiome meta-analysis’, Nature methods, 15: 796–98.

Govindarajan, K., J. MacSharry, P. G. Casey, F. Shanahan, S. A. Joyce, and C. G. Gahan. 2016. ‘Unconjugated Bile Acids Influence Expression of Circadian Genes: A Potential Mechanism for Microbe-Host Crosstalk’, PloS one, 11: e0167319.

Greenwood, B. N., R. S. Thompson, M. R. Opp, and M. Fleshner. 2014. ‘Repeated exposure to conditioned fear stress increases anxiety and delays sleep recovery following exposure to an acute traumatic stressor’, Frontiers in Psychiatry, 5: 146.

Gu, J., J. M. Thomas-Ahner, K. M. Riedl, M. T. Bailey, Y. Vodovotz, S. J. Schwartz, and S. K. Clinton. 2019. ‘Dietary Black Raspberries Impact the Colonic Microbiome and Phytochemical Metabolites in Mice’, Molecular nutrition & food research, 63: e1800636.

Haro, C., S. Garcia-Carpintero, J. F. Alcala-Diaz, F. Gomez-Delgado, J. Delgado-Lista, P. Perez-Martinez, O. A. Rangel Zuniga, G. M. Quintana-Navarro, B. B. Landa, J. C. Clemente, J. Lopez-Miranda, A. Camargo, and F. Perez-Jimenez. 2016. ‘The gut microbial community in metabolic syndrome patients is modified by diet’, J Nutr Biochem, 27: 27–31.

Herfel, T. M., S. K. Jacobi, X. Lin, V. Fellner, D. C. Walker, Z. E. Jouni, and J. Odle. 2011. ‘Polydextrose enrichment of infant formula demonstrates prebiotic characteristics by altering intestinal microbiota, organic acid concentrations, and cytokine expression in suckling piglets’, The Journal of nutrition, 141: 2139–45.

Hillmann, B., G. A. Al-Ghalith, R. R. Shields-Cutler, Q. Zhu, D. M. Gohl, K. B. Beckman, R. Knight, and D. Knights. 2018. ‘Evaluating the Information Content of Shallow Shotgun Metagenomics’, mSystems, 3.

Ho, K. J. 1976. ‘Circadian distribution of bile acids in the enterohepatic circulatory system in rats’, The American journal of physiology, 230: 1331–5.

Huang, C., J. Chen, J. Wang, H. Zhou, Y. Lu, L. Lou, J. Zheng, L. Tian, X. Wang, Z. Cao, and Y. Zeng. 2017. ‘Dysbiosis of Intestinal Microbiota and Decreased Antimicrobial Peptide Level in Paneth Cells during Hypertriglyceridemia-Related Acute Necrotizing Pancreatitis in Rats’, Front Microbiol, 8: 776.

Human Microbiome Project, Consortium. 2012. ‘Structure, function and diversity of the healthy human microbiome’, Nature, 486: 207–14.

Janssen, S., D. McDonald, A. Gonzalez, J. A. Navas-Molina, L. Jiang, Z. Z. Xu, K. Winker, D. M. Kado, E. Orwoll, M. Manary, S. Mirarab, and R. Knight. 2018. ‘Phylogenetic Placement of Exact Amplicon Sequences Improves Associations with Clinical Information’, mSystems, 3.

Kaczmarek, A., K. Skowron, A. Budzynska, and E. Gospodarek-Komkowska. 2018. ‘Virulence-associated genes and antibiotic susceptibility among vaginal and rectal Escherichia coli isolates from healthy pregnant women in Poland’, Folia Microbiol (Praha), 63: 637–43.

Kelly, B. J., R. Gross, K. Bittinger, S. Sherrill-Mix, J. D. Lewis, R. G. Collman, F. D. Bushman, and H. Li. 2015. ‘Power and sample-size estimation for microbiome studies using pairwise distances and PERMANOVA’, Bioinformatics, 31: 2461–8.

Kervezee, L., A. Kosmadopoulos, and D. B. Boivin. 2020. ‘Metabolic and cardiovascular consequences of shift work: The role of circadian disruption and sleep disturbances’, The European journal of neuroscience, 51: 396–412.

Keskey, R., E. Papazian, A. Lam, T. Toni, S. Hyoju, R. Thewissen, A. Zaborin, O. Zaborina, and J. C. Alverdy. 2020. ‘Defining Microbiome Readiness for Surgery: Dietary Prehabilitation and Stool Biomarkers as Predictive Tools to Improve Outcome’, Ann Surg.

Klosterbuer, A., Z. F. Roughead, and J. Slavin. 2011. ‘Benefits of dietary fiber in clinical nutrition’, Nutr Clin Pract, 26: 625–35.

Knights, D., J. Kuczynski, O. Koren, R. E. Ley, D. Field, R. Knight, T. Z. DeSantis, and S. T. Kelley. 2011. ‘Supervised classification of microbiota mitigates mislabeling errors’, The ISME journal, 5: 570–3.

Koecher, K. J., W. Thomas, and J. L. Slavin. 2015. ‘Healthy subjects experience bowel changes on enteral diets: addition of a fiber blend attenuates stool weight and gut bacteria decreases without changes in gas’, JPEN. Journal of parenteral and enteral nutrition, 39: 337–43.

Koh, G. Y., A. Kane, X. Wu, J. Mason, and J. Crott. 2019. ‘Parabacteroides Distasonis Attenuates Tumorigenesis, Modulates Inflammatory Markers, and Promotes Intestinal Barrier Integrity in Azoxymethane-treated Mice (OR04-02-19)’, Curr Dev Nutr, 3.

Kohsaka, A., A. D. Laposky, K. M. Ramsey, C. Estrada, C. Joshu, Y. Kobayashi, F. W. Turek, and J. Bass. 2007. ‘High-fat diet disrupts behavioral and molecular circadian rhythms in mice’, Cell metabolism, 6: 414–21.

Leone, V., S. M. Gibbons, K. Martinez, A. L. Hutchison, E. Y. Huang, C. M. Cham, J. F. Pierre, A. F. Heneghan, A. Nadimpalli, N. Hubert, E. Zale, Y. Wang, Y. Huang, B. Theriault, A. R. Dinner, M. W. Musch, K. A. Kudsk, B. J. Prendergast, J. A. Gilbert, and E. B. Chang. 2015. ‘Effects of diurnal variation of gut microbes and high-fat feeding on host circadian clock function and metabolism’, Cell host & microbe, 17: 681–9.

Liu, S., P. Qin, and J. Wang. 2019. ‘High-Fat Diet Alters the Intestinal Microbiota in Streptozotocin-Induced Type 2 Diabetic Mice’, Microorganisms, 7.

Lozupone, C., and R. Knight. 2005. ‘UniFrac: a new phylogenetic method for comparing microbial communities’, Applied and environmental microbiology, 71: 8228–35.

Lundasen, T., C. Galman, B. Angelin, and M. Rudling. 2006. ‘Circulating intestinal fibroblast growth factor 19 has a pronounced diurnal variation and modulates hepatic bile acid synthesis in man’, Journal of internal medicine, 260: 530–6.

Ma, K., R. Xiao, H. T. Tseng, L. Shan, L. Fu, and D. D. Moore. 2009. ‘Circadian dysregulation disrupts bile acid homeostasis’, PloS one, 4: e6843.

Maki, K. A., L. A. Burke, M. W. Calik, M. Watanabe-Chailland, D. Sweeney, L. E. Romick-Rosendale, S. J. Green, and A. M. Fink. 2020. ‘Sleep fragmentation increases blood pressure and is associated with alterations in the gut microbiome and fecal metabolome in rats’, Physiological genomics, 52: 280–92.

Maltz, R. M., J. Keirsey, S. C. Kim, A. R. Mackos, R. Z. Gharaibeh, C. C. Moore, J. Xu, A. Somogyi, and M. T. Bailey. 2019. ‘Social Stress Affects Colonic Inflammation, the Gut Microbiome, and Short-chain Fatty Acid Levels and Receptors’, Journal of pediatric gastroenterology and nutrition, 68: 533–40.

Mandal, S., W. Van Treuren, R. A. White, M. Eggesbo, R. Knight, and S. D. Peddada. 2015. ‘Analysis of composition of microbiomes: a novel method for studying microbial composition’, Microb Ecol Health Dis, 26: 27663.

Marcheva, B., K. M. Ramsey, E. D. Buhr, Y. Kobayashi, H. Su, C. H. Ko, G. Ivanova, C. Omura, S. Mo, M. H. Vitaterna, J. P. Lopez, L. H. Philipson, C. A. Bradfield, S. D. Crosby, L. JeBailey, X. Wang, J. S. Takahashi, and J. Bass. 2010. ‘Disruption of the clock components CLOCK and BMAL1 leads to hypoinsulinaemia and diabetes’, Nature, 466: 627–31.

Marco, M. L., M. E. Sanders, M. Ganzle, M. C. Arrieta, P. D. Cotter, L. De Vuyst, C. Hill, W. Holzapfel, S. Lebeer, D. Merenstein, G. Reid, B. E. Wolfe, and R. Hutkins. 2021. ‘The International Scientific Association for Probiotics and Prebiotics (ISAPP) consensus statement on fermented foods’, Nat Rev Gastroenterol Hepatol.

Maslanik, T., L. Mahaffey, K. Tannura, L. Beninson, B. N. Greenwood, and M. Fleshner. 2013. ‘The inflammasome and danger associated molecular patterns (DAMPs) are implicated in cytokine and chemokine responses following stressor exposure’, Brain, behavior, and immunity, 28: 54–62.

McDonald, D., M. N. Price, J. Goodrich, E. P. Nawrocki, T. Z. DeSantis, A. Probst, G. L. Andersen, R. Knight, and P. Hugenholtz. 2012. ‘An improved Greengenes taxonomy with explicit ranks for ecological and evolutionary analyses of bacteria and archaea’, The ISME journal, 6: 610–8.

McMillin, M., and S. DeMorrow. 2016. ‘Effects of bile acids on neurological function and disease’, FASEB J, 30: 3658–68.

Mekhjian, H. S., S. F. Phillips, and A. F. Hofmann. 1979. ‘Colonic absorption of unconjugated bile acids: perfusion studies in man’, Digestive diseases and sciences, 24: 545–50.

Melnik, A. V., R. R. da Silva, E. R. Hyde, A. A. Aksenov, F. Vargas, A. Bouslimani, I. Protsyuk, A. K. Jarmusch, A. Tripathi, T. Alexandrov, R. Knight, and P. C. Dorrestein. 2017. ‘Coupling Targeted and Untargeted Mass Spectrometry for Metabolome-Microbiome-Wide Association Studies of Human Fecal Samples’, Analytical chemistry, 89: 7549–59.

Mertens, K. L., A. Kalsbeek, M. R. Soeters, and H. M. Eggink. 2017. ‘Bile Acid Signaling Pathways from the Enterohepatic Circulation to the Central Nervous System’, Frontiers in neuroscience, 11: 617.

Mika, A., H. E. Day, A. Martinez, N. L. Rumian, B. N. Greenwood, M. Chichlowski, B. M. Berg, and M. Fleshner. 2016. ‘Early life diets with prebiotics and bioactive milk fractions attenuate the impact of stress on learned helplessness behaviours and alter gene expression within neural circuits important for stress resistance’, The European journal of neuroscience.

Mika, A., and M. Fleshner. 2016. ‘Early-life exercise may promote lasting brain and metabolic health through gut bacterial metabolites’, Immunology and cell biology, 94: 151–7.

Mika, A., M. Gaffney, R. Roller, A. Hills, C. A. Bouchet, K. A. Hulen, R. S. Thompson, M. Chichlowski, B. M. Berg, and M. Fleshner. 2018. ‘Feeding the developing brain: Juvenile rats fed diet rich in prebiotics and bioactive milk fractions exhibit reduced anxiety-related behavior and modified gene expression in emotion circuits’, Neuroscience letters.

Morris, C. J., T. E. Purvis, K. Hu, and F. A. Scheer. 2016. ‘Circadian misalignment increases cardiovascular disease risk factors in humans’, Proceedings of the National Academy of Sciences of the United States of America, 113: E1402–11.

Olivadoti, M. D., and M. R. Opp. 2008. ‘Effects of i.c.v. administration of interleukin-1 on sleep and body temperature of interleukin-6-deficient mice’, Neuroscience, 153: 338–48.

Oosterman, J. E., A. Kalsbeek, S. E. la Fleur, and D. D. Belsham. 2015. ‘Impact of nutrients on circadian rhythmicity’, American journal of physiology. Regulatory, integrative and comparative physiology, 308: R337–50.

Ovacik, M. A., S. Sukumaran, R. R. Almon, D. C. DuBois, W. J. Jusko, and I. P. Androulakis. 2010. ‘Circadian signatures in rat liver: from gene expression to pathways’, BMC bioinformatics, 11: 540.

Perino, A., H. Demagny, L. A. Velazquez-Villegas, and K. Schoonjans. 2020. ‘Molecular Physiology of Bile Acid Signaling in Health, Disease and Aging’, Physiological reviews.

Saha, D. C., and R. A. Reimer. 2014. ‘Long-term intake of a high prebiotic fiber diet but not high protein reduces metabolic risk after a high fat challenge and uniquely alters gut microbiota and hepatic gene expression’, Nutrition research, 34: 789–96.

Saulnier, D. M., Y. Ringel, M. B. Heyman, J. A. Foster, P. Bercik, R. J. Shulman, J. Versalovic, E. F. Verdu, T. G. Dinan, G. Hecht, and F. Guarner. 2013. ‘The intestinal microbiome, probiotics and prebiotics in neurogastroenterology’, Gut microbes, 4: 17–27.

Sepe, V., C. Festa, B. Renga, A. Carino, S. Cipriani, C. Finamore, D. Masullo, F. Del Gaudio, M. C. Monti, S. Fiorucci, and A. Zampella. 2016. ‘Insights on FXR selective modulation. Speculation on bile acid chemical space in the discovery of potent and selective agonists’, Scientific reports, 6: 19008.

Shapiro, J., N. A. Cohen, V. Shalev, A. Uzan, O. Koren, and N. Maharshak. 2019. ‘Psoriatic patients have a distinct structural and functional fecal microbiota compared with controls’, J Dermatol, 46: 595–603.

Silvennoinen, R., H. Quesada, I. Kareinen, J. Julve, L. Kaipiainen, H. Gylling, F. Blanco-Vaca, J. C. Escola-Gil, P. T. Kovanen, and M. Lee-Rueckert. 2015. ‘Chronic intermittent psychological stress promotes macrophage reverse cholesterol transport by impairing bile acid absorption in mice’, Physiological reports, 3.

Staley, C., A. R. Weingarden, A. Khoruts, and M. J. Sadowsky. 2017. ‘Interaction of gut microbiota with bile acid metabolism and its influence on disease states’, Applied microbiology and biotechnology, 101: 47–64.

Tahara, Y., M. Yamazaki, H. Sukigara, H. Motohashi, H. Sasaki, H. Miyakawa, A. Haraguchi, Y. Ikeda, S. Fukuda, and S. Shibata. 2018. ‘Gut Microbiota-Derived Short Chain Fatty Acids Induce Circadian Clock Entrainment in Mouse Peripheral Tissue’, Scientific reports, 8: 1395.

Tang, R., Y. Jiang, A. Tan, J. Ye, X. Xian, Y. Xie, Q. Wang, Z. Yao, and Z. Mo. 2018. ‘16S rRNA gene sequencing reveals altered composition of gut microbiota in individuals with kidney stones’, Urolithiasis, 46: 503–14.

Tang, Z. Z., G. Chen, and A. V. Alekseyenko. 2016. ‘PERMANOVA-S: association test for microbial community composition that accommodates confounders and multiple distances’, Bioinformatics, 32: 2618–25.

Tarr, A. J., J. D. Galley, S. E. Fisher, M. Chichlowski, B. M. Berg, and M. T. Bailey. 2015. ’The prebiotics 3’Sialyllactose and 6’Sialyllactose diminish stressor-induced anxiety-like behavior and colonic microbiota alterations: Evidence for effects on the gut-brain axis’, Brain, behavior, and immunity, 50: 166–77.

Thaiss, C. A., D. Zeevi, M. Levy, G. Zilberman-Schapira, J. Suez, A. C. Tengeler, L. Abramson, M. N. Katz, T. Korem, N. Zmora, Y. Kuperman, I. Biton, S. Gilad, A. Harmelin, H. Shapiro, Z. Halpern, E. Segal, and E. Elinav. 2014. ‘Transkingdom control of microbiota diurnal oscillations promotes metabolic homeostasis’, Cell, 159: 514–29.

Thomas, C., R. Pellicciari, M. Pruzanski, J. Auwerx, and K. Schoonjans. 2008. ‘Targeting bile-acid signalling for metabolic diseases’, Nat Rev Drug Discov, 7: 678–93.

Thompson, R. S., J. P. Christianson, T. M. Maslanik, S. F. Maier, B. N. Greenwood, and M. Fleshner. 2013. ‘Effects of stressor controllability on diurnal physiological rhythms’, Physiology & behavior, 112-113: 32–39.

Thompson, R. S., R. Roller, B. N. Greenwood, and M. Fleshner. 2016. ‘Wheel running improves REM sleep and attenuates stress-induced flattening of diurnal rhythms in F344 rats’, Stress: 1–13.

Thompson, R. S., R. Roller, A. Mika, B. N. Greenwood, R. Knight, M. Chichlowski, B. M. Berg, and M. Fleshner. 2017. ‘Dietary Prebiotics and Bioactive Milk Fractions Improve NREM Sleep, Enhance REM Sleep Rebound and Attenuate the Stress-Induced Decrease in Diurnal Temperature and Gut Microbial Alpha Diversity’, Front Behav Neurosci, 10: 240.

Thompson, R. S., P. V. Strong, P. J. Clark, T. M. Maslanik, K. P. Wright, Jr., B. N. Greenwood, and M. Fleshner. 2014. ‘Repeated fear-induced diurnal rhythm disruptions predict PTSD-like sensitized physiological acute stress responses in F344 rats’, Acta physiologica.

Thompson, R. S., F. Vargas, P. C. Dorrestein, M. Chichlowski, B. M. Berg, and M. Fleshner. 2020. ‘Dietary prebiotics alter novel microbial dependent fecal metabolites that improve sleep’, Scientific reports, 10: 3848.

Tognini, P., M. Murakami, Y. Liu, K. L. Eckel-Mahan, J. C. Newman, E. Verdin, P. Baldi, and P. Sassone-Corsi. 2017. ‘Distinct Circadian Signatures in Liver and Gut Clocks Revealed by Ketogenic Diet’, Cell metabolism, 26: 523–38 e5.

Turek, F. W., C. Joshu, A. Kohsaka, E. Lin, G. Ivanova, E. McDearmon, A. Laposky, S. Losee-Olson, A. Easton, D. R. Jensen, R. H. Eckel, J. S. Takahashi, and J. Bass. 2005. ‘Obesity and metabolic syndrome in circadian Clock mutant mice’, Science, 308: 1043–5.

van Meer, H., G. Boehm, F. Stellaard, A. Vriesema, J. Knol, R. Havinga, P. J. Sauer, and H. J. Verkade. 2008. ‘Prebiotic oligosaccharides and the enterohepatic circulation of bile salts in rats’, American journal of physiology. Gastrointestinal and liver physiology, 294: G540–7.

Vitaterna, M. H., K. Shimomura, and P. Jiang. 2019. ‘Genetics of Circadian Rhythms’, Neurol Clin, 37: 487–504.

Voigt, R. M., C. B. Forsyth, S. J. Green, P. A. Engen, and A. Keshavarzian. 2016. ‘Circadian Rhythm and the Gut Microbiome’, International review of neurobiology, 131: 193–205.

Wilson, K. H. 1983. ‘Efficiency of various bile salt preparations for stimulation of Clostridium difficile spore germination’, Journal of clinical microbiology, 18: 1017–9.

Xie, X., Y. He, H. Li, D. Yu, L. Na, T. Sun, D. Zhang, X. Shi, Y. Xia, T. Jiang, S. Rong, S. Yang, X. Ma, and G. Xu. 2018. ‘Effects of prebiotics on immunologic indicators and intestinal microbiota structure in perioperative colorectal cancer patients’, Nutrition, 61: 132–42.

Yatsunenko, T., F. E. Rey, M. J. Manary, I. Trehan, M. G. Dominguez-Bello, M. Contreras, M. Magris, G. Hidalgo, R. N. Baldassano, A. P. Anokhin, A. C. Heath, B. Warner, J. Reeder, J. Kuczynski, J. G. Caporaso, C. A. Lozupone, C. Lauber, J. C. Clemente, D. Knights, R. Knight, and J. I. Gordon. 2012. ‘Human gut microbiome viewed across age and geography’, Nature, 486: 222–7.

Yilmaz, P., L. W. Parfrey, P. Yarza, J. Gerken, E. Pruesse, C. Quast, T. Schweer, J. Peplies, W. Ludwig, and F. O. Glockner. 2014. ‘The SILVA and “All-species Living Tree Project (LTP)” taxonomic frameworks’, Nucleic acids research, 42: D643–8.

Yu, M., H. Jia, C. Zhou, Y. Yang, Y. Zhao, M. Yang, and Z. Zou. 2017. ‘Variations in gut microbiota and fecal metabolic phenotype associated with depression by 16S rRNA gene sequencing and LC/MS-based metabolomics’, J Pharm Biomed Anal, 138: 231–39.

Yu, Z., J. Yang, D. Xiang, G. Li, D. Liu, and C. Zhang. 2020. ‘Circadian rhythms and bile acid homeostasis: a comprehensive review’, Chronobiology international: 1–11.

Yutin, N., and M. Y. Galperin. 2013. ‘A genomic update on clostridial phylogeny: Gram-negative spore formers and other misplaced clostridia’, Environ Microbiol, 15: 2631–41.

Zheng, S. N., S. S. Zhang, M. Y. Yu, J. Tang, X. M. Lu, F. Wang, J. Y. Yang, and F. M. Li. 2011. ‘An H-1 NMR and UPLC-MS-based plasma metabonomic study to investigate the biochemical changes in chronic unpredictable mild stress model of depression’, Metabolomics, 7: 413–23.

